# Maternal IL-10 restricts fetal emergency myelopoiesis

**DOI:** 10.1101/2023.09.13.557548

**Authors:** Amélie Collins, James W. Swann, Melissa A. Proven, Chandani M. Patel, Carl A. Mitchell, Monica Kasbekar, Paul V. Dellorusso, Emmanuelle Passegué

## Abstract

Neonates, in contrast to adults, are highly susceptible to inflammation and infection. Here we investigate how late fetal liver (FL) mouse hematopoietic stem and progenitor cells (HSPC) respond to inflammation, testing the hypothesis that deficits in engagement of emergency myelopoiesis (EM) pathways limit neutrophil output and contribute to perinatal neutropenia. We show that despite similar molecular wiring as adults, fetal HSPCs have limited production of myeloid cells at steady state and fail to activate a classical EM transcriptional program. Moreover, we find that fetal HSPCs are capable of responding to EM-inducing inflammatory stimuli *in vitro*, but are restricted by maternal anti-inflammatory factors, primarily interleukin-10 (IL-10), from activating EM pathways *in utero*. Accordingly, we demonstrate that loss of maternal IL-10 restores EM activation in fetal HSPCs but at the cost of premature parturition. These results reveal the evolutionary trade-off inherent in maternal anti-inflammatory responses that maintain pregnancy but render the fetus unresponsive to EM activation signals and susceptible to infection.

**HIGHLIGHTS:**

- The structure of the HSPC compartment is conserved from late fetal to adult life.
- Fetal HSPCs have diminished steady-state myeloid cell production compared to adult.
- Fetal HSPCs are restricted from engaging in emergency myelopoiesis by maternal IL-10.
- Restriction of emergency myelopoiesis may explain neutropenia in septic neonates.

**eTOC BLURB:** Fetal hematopoietic stem and progenitor cells are restricted from activating emergency myelopoiesis pathways by maternal IL-10, resulting in inadequate myeloid cell production in response to inflammatory challenges and contributing to neonatal neutropenia.

## INTRODUCTION

Self-renewing hematopoietic stem cells (HSC) sit atop the hematopoietic hierarchy and control blood production both at homeostasis and in response to diverse demands across the lifetime of organisms^1^. HSCs differentiate through a series of intermediary lineage-biased multipotent progenitors (MPP)^2,3^ and a range of lineage-committed myeloid or lymphoid progenitor cells, to give rise to all innate and adaptive immune cells that protect the organism from invading pathogens. Demand-adapted production from these hematopoietic stem and progenitor cells (HSPC) is an essential feature of the hematopoietic response to systemic inflammation^4^. Each period of the lifespan (e.g., fetal, childhood, adulthood, aging) also superimposes features that modulate hematopoiesis to affect immune responses^5^, with the perinatal period posing distinct immunological challenges^6^. Maternal tolerance for the semi-allogeneic fetus is essential for continued pregnancy, which can be threatened by inflammation. Postnatally, developing immune memory to pathogen exposure must be balanced with tolerance to commensal microbes and food antigens. Inflammatory responses are therefore tightly regulated by numerous anti-inflammatory cytokines and mechanisms. Occurring in the context of this anti-inflammatory milieu^7^, infections in the perinatal period cause high morbidity and mortality, and neonatal sepsis is a significant global health burden^8^. A well-recognized feature of the neonatal response to infection that contributes to these poor outcomes is neutropenia caused by inadequate neutrophil production^9,10^. In both human and rodent experimental models, neonatal neutropenia has been attributed to a failure of myeloid progenitor cells to meet demand^11–13^, but the mechanisms underlying the deficient production of myeloid cells in neonates remain largely unknown.

During fetal development, definitive self-renewing HSCs emerge by embryonic day 10 (E10) in mice^14^ and 5 weeks post-conception in humans^15^, long before the perinatal period. Fetal HSCs are more proliferative, have higher self-renewal potential, and have different lineage output than adult HSCs^16,17^, with perinatal blood production dominated by erythropoiesis and myelopoiesis, and limited lymphopoiesis. While many of the lineage-specific transcription factors are shared across the ages, fetal HSCs are governed by distinct molecular regulators including Sox17^18^, CEBPα^19^, and the Lin28b-let7-Hmga2 axis^20^. Until recently, the prevailing model of embryonic hematopoiesis developed in murine models was that upon emergence from endothelial precursors in the aorta-gonad-mesonephros (AGM) region of the dorsal aorta, HSCs migrate to the fetal liver (FL), where they divide rapidly, meeting the heightened need for mature blood cells in the developing embryo and generating a pool of cells that become adult HSCs as they progressively transition from the FL to the bone marrow (BM) cavity in the perinatal period. Several recent studies have challenged some of these assumptions. First, phenotypically-defined fetal HSCs have been found to contribute minimally to blood production during early life, which is instead largely supported by an HSC-independent MPP pool^21,22^. A comparison of HSCs and MPPs in FL vs. fetal BM also concluded that while HSPCs are detected in the BM during late fetal timepoints (E15.5 – P0), they are markedly reduced in number and largely non-functional, only acquiring robust stem-like function post-natally^23^. Second, while FL HSCs do indeed divide frequently, correlating with their high “repopulating activity” in transplantation assays, their contribution to the adult HSC pool as measured by lineage tracking approaches appears to be minimal^21, 24^, with most adult HSCs generated instead via post-natal expansion in the BM^24^. Collectively, the revised model of perinatal hematopoiesis emerging from these studies suggests that fetal HSCs act mainly as a backup reservoir, with most functional blood production at steady state deriving from FL MPPs. However, with the exception of the megakaryocytic/erythroid-biased MPP2^23^, the functional output of the fetal MPP compartment has not been investigated and it remains unclear how this developing framework of fetal hematopoiesis functions in the context of clinically relevant challenges of perinatal life such as sepsis and neutropenia.

In adults, the process by which the hematopoietic system rapidly generates myeloid cells in response to inflammation is called emergency myelopoiesis (EM)^25,26^. EM functions at multiple levels of the adult HSPC hierarchy, all geared towards rapid amplification of myeloid cell production, including neutrophils, followed by restoration of homeostatic levels. In response to pro-inflammatory cytokines and growth factors, HSCs are activated out of their quiescent state and initiate a cascade of events that amplifies myelopoiesis at the expense of other blood lineages^27–33^. At the level of the MPP compartment in mice, EM activation redirects lymphoid-biased MPP4 toward the myeloid lineage^34^ and over-produces myeloid-biased MPP3^2,35^, with the specific expansion of a self-reinforcing secretory FcγR^+^ MPP3 subset^36^, driving the formation of GMP clusters in the BM cavity^37^, which serve as hubs to rapidly scale-up neutrophil production^4^. Currently, little is known about the ability of fetal HSPCs to respond to inflammation and activate classical adult-like EM pathways. Here, we set out to investigate how late FL mouse HSPCs (E18.5) react to inflammatory stimulus both *ex vivo* in culture conditions and *in vivo* in pregnant dams.

## RESULTS

### Conserved lineage-priming in fetal and neonatal HSPCs

To investigate the structure of the early hematopoietic cell hierarchy in fetal and neonatal timepoints, we used the phenotypic definition of HSPCs previously established in adult mice in the Lin^-^/Sca-1^+^/c-Kit^+^ (LSK) BM fraction (**Figure S1A**)^2^. HSCs were identified as Flk2^-^/CD150^+^/CD48^-^ LSK, MPP2 as Flk2^-^/CD150^+^/CD48^+^ LSK, MPP3 as Flk2^-^/CD150^-^/CD48^+^ LSK, and MPP4 as Flk2^+^/CD150^-^ LSK in the BM, spleen, and liver during the neonatal period (P1 to P21) and in the FL during late embryonic development (E12.5 to E18.5). Of note, we excluded Mac-1 and CD4 from the lineage cocktail in the fetal timepoints due to their known expression on fetal HSCs^38^. As already described^23,24,38,39^, we found phenotypically defined HSCs in the FL as early as E12.5 and growing in number throughout embryonic development (**Figure 1A**). After birth, the liver was rapidly depleted of HSCs, with increasing numbers seen in the spleen and BM in neonates. HSCs persisted in neonatal spleen until ∼ 2 weeks of age, after which they were found primarily in the BM. Phenotypically defined MPPs could also be identified as early as E12.5, with numbers expanding in the FL during development and migrating through the spleen to the BM in neonates with kinetics similar to that of HSCs (**Figure S1B**). In both cases, HSC and MPP numbers did not reach adult levels until ∼ 6 weeks of age. To confirm the lineage priming of fetal/neonatal HSPCs, we chose to focus on one FL timepoint (E18.5) and one neonatal BM timepoint (P14) in comparison to adult BM obtained from 8-week-old mice (**Figure 1B**). We first performed bulk RNA sequencing (RNA-seq) on HSCs, MPP2s, MPP3s and MPP4s isolated from each of these developmental stages. Principal component (PC) analysis demonstrated that the different HSPC subsets were transcriptionally similar across development, with the type of HSPCs (HSC, MPP2, MPP3 or MPP4) explaining more variability in gene expression in PC1 than their developmental origin (fetal, neonatal, or adult) in PC2 (**Figure 1C**). Differentially expressed gene (DEG) analyses comparing MPP2, MPP3, and MPP4 to HSC at each developmental stage also found many shared DEGs, with pathway analysis highlighting the expected megakaryocyte/erythrocyte bias in MPP2, myeloid bias in MPP3, and lymphoid bias in MPP4 (**Figure 1D**)^2^. These results reinforce the idea that the HSPC hierarchy is conserved across development^23^ and show that lineage biased MPP subsets can be isolated at fetal and neonatal timepoints using similar surface markers as in adults.

**Figure 1.**
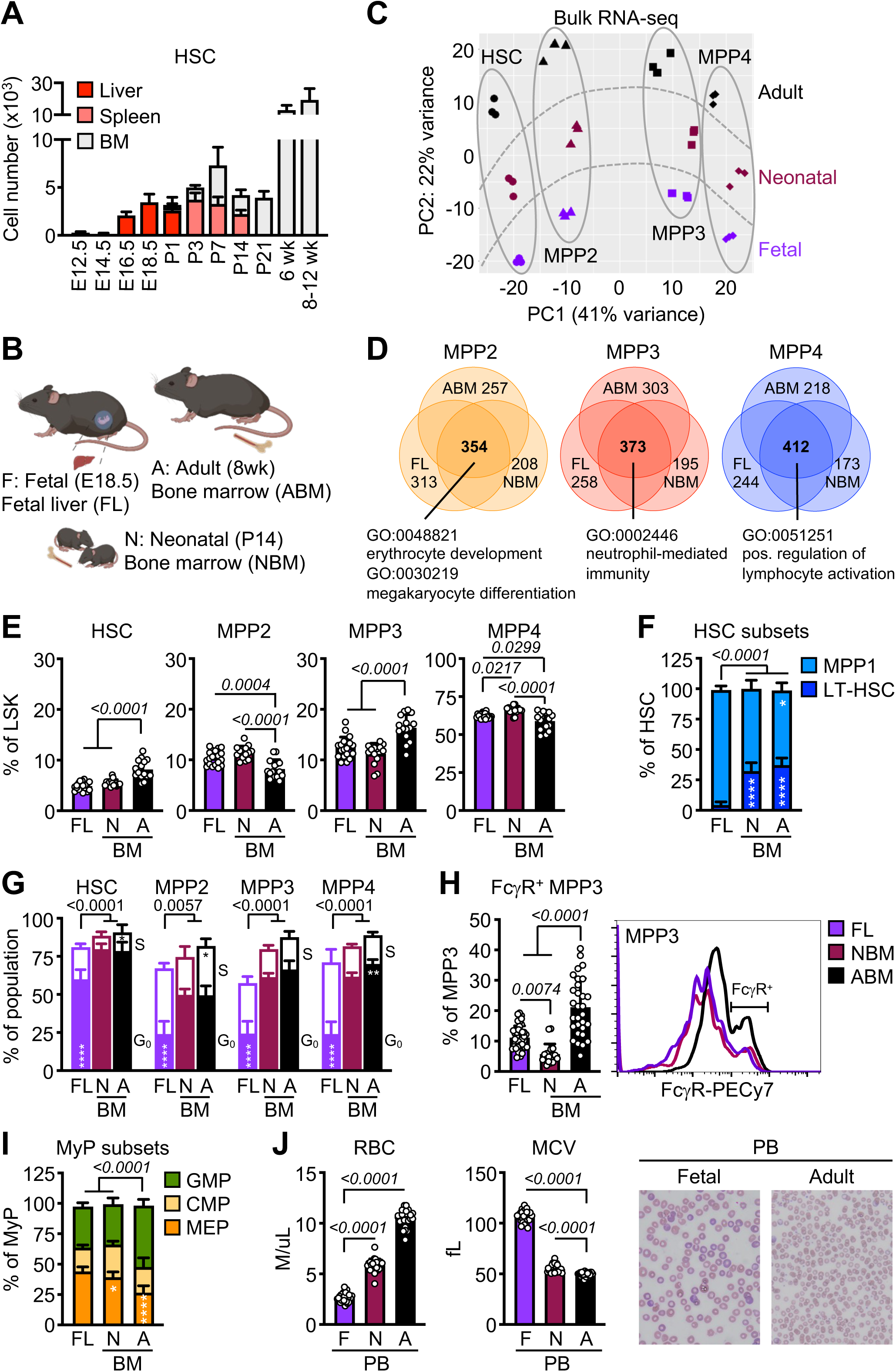
Conserved lineage-priming in fetal and neonatal HSPCs. **A)** HSC number in liver, spleen, or bone marrow (BM) of fetuses (F): E12.5 (6), E14.5 (13), E16.5 (7), and E18.5 (8); neonates (N): P1 (4), P3 (4), P7 (9), P14 (5), and P21 (5); and adult (A) mice: 6wk (6) and 8-12wk (7); wk, week. Results are from the number of biological repeat (individual mice) from at least 2 independent experiments. **B)** Schematic of selected fetal liver (FL), neonatal BM (NBM), and adult BM (ABM) timepoints. **C)** Principal component analysis of bulk RNA-seq of FL, NBM, and ABM HSPC populations. Results are from 3 independent sets of pooled fetuses, neonate, and adult mice, with all HSPC populations in a given replicate coming from the same pool. **D)** Venn diagrams showing the number of unique and shared differentially enriched genes (DEG, log_2_ FC > 1, FDR < 0.05) found at all ages in each MPP population compared to HSCs. Selected gene ontology (GO) pathways found enriched in shared DEGs are shown below. **E)** Frequency of the indicated HSPC populations within FL, NBM, and ABM LSK cells (3 independent experiments). **F)** Frequency of MPP1 and long-term HSC (LT-HSC) within FL, NBM, and ABM HSCs (3 independent experiments). **G)** G_0_ and S phase distribution in HSPCs as measured in *Fucci2* cell cycle reporter mice of indicated ages (5 independent experiments). **H)** Frequency of FcγR^+^ MPP3 (left) and representative FACS plot of FcγR staining (right) on FL, NBM, and ABM MPP3 (3 independent experiments). **I)** Frequency of GMPs, CMPs, and MEPs within the FL, NBM, and ABM myeloid progenitors (MyP, Lin^-^/c-Kit^+^/Sca-1^-^) compartment (3 independent experiments). **J)** Red blood cell (RBC) and mean corpuscular volume (MCV) from complete blood cell (CBC) counts of fetal (F), neonate (N) and adult (A) peripheral blood (PB), with representative Wright-Giemsa staining of fetal and adult PB smears (5 independent experiments). Data are means ± S.D.; *p_adj_ ≤ 0.05; **p_adj_ ≤ 0.01; ****p_adj_ ≤ 0.0001. Circles represent biological replicates (individual mice). See also Figure S1.

To determine whether there was a myelopoietic deficit in fetal and neonatal HSPCs, we next investigated their differentiation properties. Consistent with the overall differences in cellularity at these development stages, fetal and neonatal mice had far fewer HSCs and MPPs compared to adult mice (**Figure S1C**). However, within the LSK compartment, fetal and neonatal mice had relatively more MPP2 and MPP4 and fewer HSCs and MPP3 than adult mice (**Figure 1E**). As expected, the HSC compartment of fetal mice was highly activated^40–42^, with fewer quiescent G_0_ fetal HSCs (**Figure 1F**), more CD34^+^ MPP1 vs. CD34^-^ long-term HSCs (**Figure 1G**), and higher Sca-1 expression (**Figure S1D**) compared to adult HSCs, with neonatal HSCs showing an intermediate transitional behavior. We also found that fetal MPPs were more highly cycling than neonatal and adult MPPs (**Figure 1G**). In contrast, adult HSCs had higher expression of CD41 than either fetal or neonatal HSCs (**Figure S1E**), a marker known to label HSCs biased for myeloid differentiation^43^. Similarly, a significantly smaller fraction of fetal and neonatal MPP3 expressed FcγR compared to adult MPP3 (**Figure 1H**), which identifies a self-reinforcing subset of MPP3 poised for accelerated myeloid differentiation^36^. In addition, we found that fetal and neonatal mice had fewer GMPs and more megakaryocyte/erythroid progenitors (MEPs) than adult mice (**Figure 1I**). Consistent with the expansion of erythroid-biased MPP2 and MEPs in fetal and neonatal mice, complete blood counts (CBC) of peripheral blood (PB) revealed fewer absolute numbers of red blood cells (RBC) but higher RBC mean corpuscular volume (MCV) than in adult blood (**Figure 1J**), indicative of active erythropoiesis. Blood smears from fetal mice also showed RBCs that were larger and more polychromatic than in adult blood, and overall fewer mature leukocytes than in neonatal and adult blood (**Figures 1J, S1F**). Altogether these results demonstrate a reduction in myeloid-biased progenitors at multiple levels of the early hematopoietic hierarchy in fetal and neonatal mice, suggesting a steady-state bias away from myelopoiesis in the perinatal period compared to adulthood.

### Fetal HSPCs have limited myeloid output

We next interrogated the ability of fetal HSPCs to produce myeloid cells. In a bulk liquid culture assay optimized to support myeloid cell growth *in vitro*^2^, fetal HSCs and MPP2 initially proliferated more rapidly than their adult counterparts but were unable to sustain the same level of myeloid cell production over time (**Figures 2A**, **S2A**). Fetal MPP3 and MPP4 also showed uniformly diminished cell proliferation and myeloid differentiation at all time points. We confirmed these findings using a myeloid/erythroid/megakaryocytic multi-lineage single cell *in vitro* differentiation assay^44^ (**Figure S2B**), observing again limited production of myeloid-containing colonies from fetal HSPCs, especially MPPs, compared to their adult counterparts (**Figures 2B, S2C**). This deficit was specific to the myeloid lineage as production of erythroid cells was uniformly increased from every fetal HSPC population compared to adult populations (**Figure 2C**). We also used an OP9 stromal cell-based version of our single cell *in vitro* differentiation assay, optimized to support lymphoid cell growth^45^, and found enhanced lymphopoiesis from fetal HSPCs with fetal MPP4s giving rise to more lymphoid-containing colonies, and lymphoid colonies from fetal HSCs and MPP4s having higher lymphoid cell numbers compared to their adult counterparts (**Figure 2D**). To further investigate functionally the lineage bias of fetal HSCs *in vivo*, we used short-term lineage tracking in sub-lethally irradiated recipient mice to measure the robustness of the myeloid output produced by 2,000 transplanted fetal or adult HSCs^2^ (**Figure 2E**). We found that despite higher chimerism, fetal HSCs quickly lose their myeloid output, with more rapid production of lymphoid cells compared to adult HSCs (**Figure 2E**). This myeloid deficit was recapitulated in the early phase of standard transplantation experiments (e.g., lethally irradiated recipients transplanted with 250 HSCs) (**Figure S2D**), with production of fewer myeloid and more lymphoid (predominantly B) cells at 1-month post-transplantation from fetal HSCs compared to adult HSCs (**Figure S2E**). However, by 2 months post-transplantation, the lineage output of fetal HSCs resembled that of adult HSCs and was maintained for over 4 months, with secondary transplantation confirming that the early myeloid deficit of fetal HSCs was not maintained over time in an adult BM environment (**Figures S2E, S2F**). Altogether, these results demonstrate that fetal HSPCs do not produce myeloid cells as robustly as adult HSPCs but instead prioritize erythropoiesis and lymphopoiesis, thereby fulfilling the hematopoietic needs of the embryonic period.

**Figure 2.**
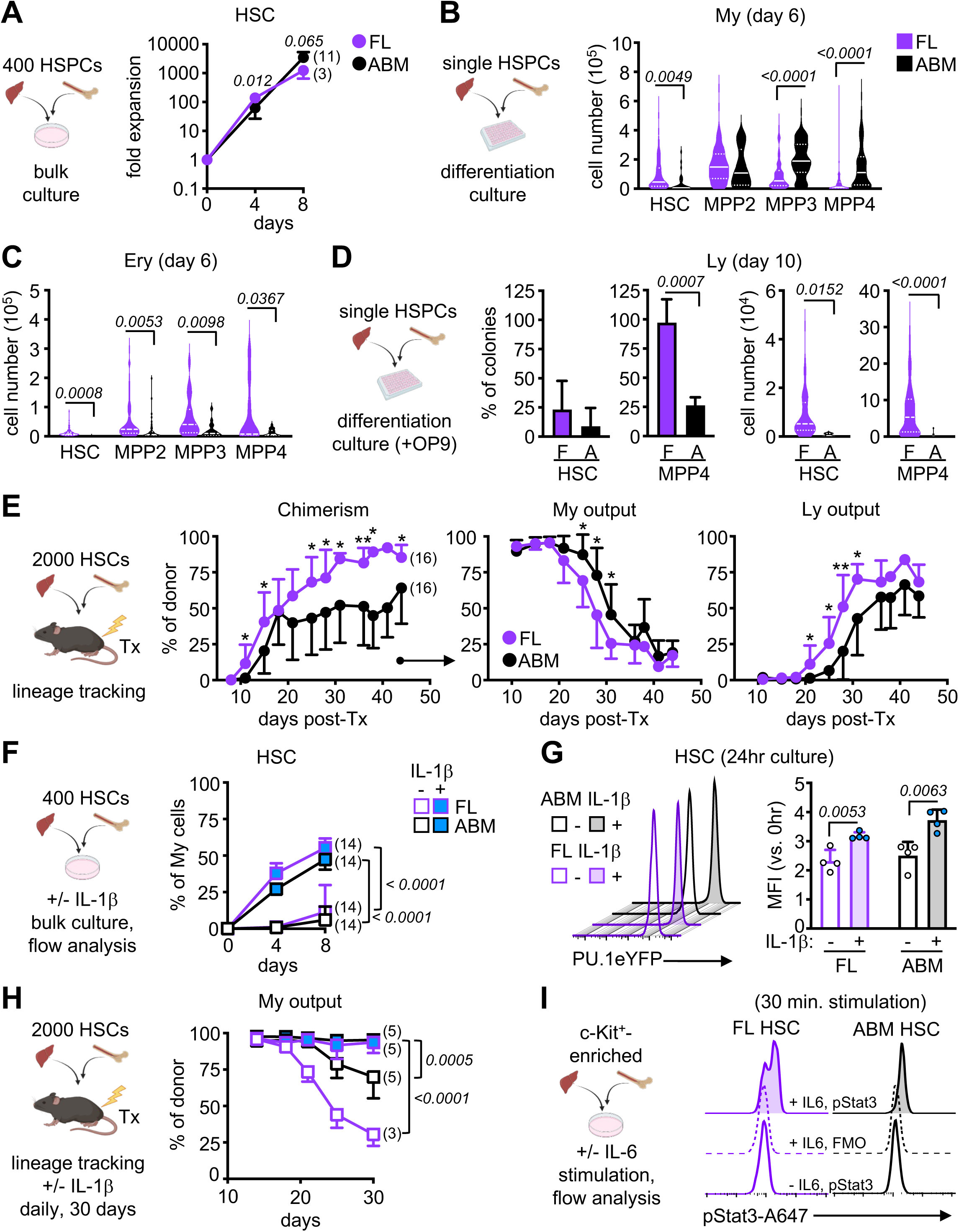
Fetal HSPCs have limited myeloid output. **A)** Schematic of bulk liquid culture and fold expansion of FL and ABM HSCs (3 independent experiments). **B-C**) Schematic of single cell differentiation culture of FL and ABM HSPCs with cellularity of myeloid (My) (B) or erythroid (Ery) (C) colonies at day 6 (5 independent experiments with a range of 11-163 colonies counted per population). Data are shown as violin plots with median and quartiles. **D)** Schematic of OP9-supported single cell differentiation culture of FL (F) and ABM (A) HSC and MPP4 (left) with proportion (middle) and cellularity (right) of lymphoid (Ly) colonies at day 10 (3 independent experiments with a range of 8-215 colonies counted per population). Cellularity data are shown as violin plots with median and quartiles. **E)** Schematic of lineage tracking transplantation (Tx) of FL and ABM HSCs in sub-lethally irradiated recipients with percent chimerism (left), and donor-derived myeloid (middle) and lymphoid (right) output in peripheral blood at the indicated days post-transplantation (4 independent experiments). **F)** Schematic of bulk liquid culture of FL or ABM HSCs with or without (+/-) IL-1β (25 ng/ml) and percent of FcγR^+^/Mac-1^+^ myeloid cells quantified by flow cytometry over time (3 independent experiments). **G)** Representative FACS histograms (left) and quantification (right) of YFP mean fluorescence intensity (MFI) of FL and ABM HSCs isolated from *PU.1-eYFP* reporter mice and cultured for 24 hours (hr) +/-IL-1β (4 independent experiments). **H)** Schematic of lineage tracking transplantation of FL and ABM HSCs in recipient mice injected daily +/-IL-1β (0.5 μg) with myeloid output in peripheral blood at the indicated times post-transplantation (1 independent experiment). **I)** Schematic of bulk liquid culture of FL or ABM HSCs +/-IL-6 (50 ng/ml) and representative FACS histograms of phospho-Stat3 fluorescence after 30 minutes (min) stimulation (2 independent experiments); FMO, fluorescence minus one. Data are means ± S.D. except for (B, C, and part of D); *p_adj_ ≤ 0.05; **p_adj_ ≤ 0.01. Number of biological replicates and transplanted mice are indicated either in parentheses or with individual circles. See also Figure S2.

### Fetal HSPCs robustly amplify myelopoiesis in response to pro-inflammatory stimuli

To directly probe the cell-intrinsic capacity of fetal HSPCs to generate myeloid-lineage cells, we assessed their response to myelopoiesis-promoting cytokines *ex vivo*. We previously demonstrated that IL-1β accelerates the production of mature myeloid cells from adult HSCs via precocious activation of PU.1, a pioneer transcription factor with a critical role in myelopoiesis^31^. Fetal HSCs grown in liquid culture in the presence of IL-1β differentiated to myeloid cells as readily as adult HSCs (**Figures 2F**, **S2G**). Using *PU.1-eYFP* reporter mice, we also showed equivalent PU.1 upregulation in fetal and adult HSCs in response to IL-1β exposure *in vitro* (**Figure 2G**). Finally, we used daily IL-1β injections to prolong the myeloid output of transplanted HSCs in sub-lethally irradiated recipients, demonstrating that fetal HSCs were as capable of producing myeloid cells as adult HSCs when exposed to a strong myeloid-promoting signal *in vivo* (**Figure 2H**). The ability of fetal HSPCs to respond to myeloid-promoting cytokines *in vitro* was not limited to IL-1β, as we also found that fetal HSPCs were equally capable as adult HSCs of phosphorylating Stat3 in response to IL-6 stimulation (**Figure 2I**), and of upregulating Sca-1 surface expression and increasing mRNA expression of interferon-stimulated genes (ISG) in response to IFNα stimulation (**Figure S2H**). Taken together, these results demonstrate that fetal HSPCs can be driven to myelopoiesis as efficiently as adult HSPCs when stimulated with pro-myeloid inflammatory factors, suggesting that fetal restriction of myelopoiesis could be attributable to decreased exposure or responsiveness to those signals.

### Transcriptional differences do not dictate the myeloid output of fetal or adult HSPCs

To determine whether fetal HSPCs harbor fundamental differences in their transcriptional landscape that might restrict their myeloid output, we performed droplet-based single cell RNA sequencing (scRNA-seq) on fetal and adult Lin^-^/c-Kit^+^ (LK) cells. Uniform manifold approximation and projection (UMAP) representation of the integrated datasets showed a common HSPC-containing center and two arms, one composed of erythroid progenitors and the other of myeloid progenitors (**Figures 3A, S3A**). Importantly, we did not identify any distinct population of cells present in one developmental stage but not in the other, reinforcing the applicability of the HSPC identification scheme across development. Cell density projection and cluster distribution recapitulated the expansion of HSCs and myeloid progenitors in the adult, and expansion of erythroid progenitors in the fetus (**Figure 3B**). We also confirmed the increase in cycling HSCs in the fetus (**Figure 3C**) and found similar numbers of genes enriched in each HSPC population when comparing DEGs between fetal and adult development stages (**Figure S3B**). Remarkably, in both scRNA-seq (**Figure 3D**) and bulk RNA-seq (**Figure 3E**) datasets, pathway analyses showed fetal HSPCs to be enriched in interferon stimulated gene (ISG) signature and adult HSPCs in antigen (Ag) processing and presentation pathways. This was already reported by others^46–48^, and validated here by qRT-PCR and surface staining (**Figures S3C, S3D**). However, we did not find compelling evidence for the contribution of either fetal or adult transcriptional signature to steady state myeloid output. While the fetal ISG signature was shown to be driven by a transient pulse of type I interferons signaling through type I interferon receptor (IFNAR) during late development and abrogated by fetal *Ifnar* deletion^46^, we found no difference in the ability of WT or *Ifnar*^-/-^ fetal HSPCs to produce myeloid or erythroid cells as measured in single cell *in vitro* differentiation assays (**Figure S3E**). This is consistent with a recent report of unchanged blood production in *Ifnar*^-/-^ neonatal mice^49^. Similarly, in adult mice, we found that loss of neither MHC class I (MHCI) nor MHC class II (MHCII) altered HSC distribution *in vivo* (**Figure S3F**), changed myeloid cell output in single cell *in vitro* differentiation assay (**Figure S3G**), or impacted short-term lineage tracking assay in sub-lethally irradiated recipients (**Figure S3H**). These results indicate that differences in the steady-state myeloid output of fetal and adult HSPCs cannot be explained by their most salient transcriptional features.

**Figure 3.**
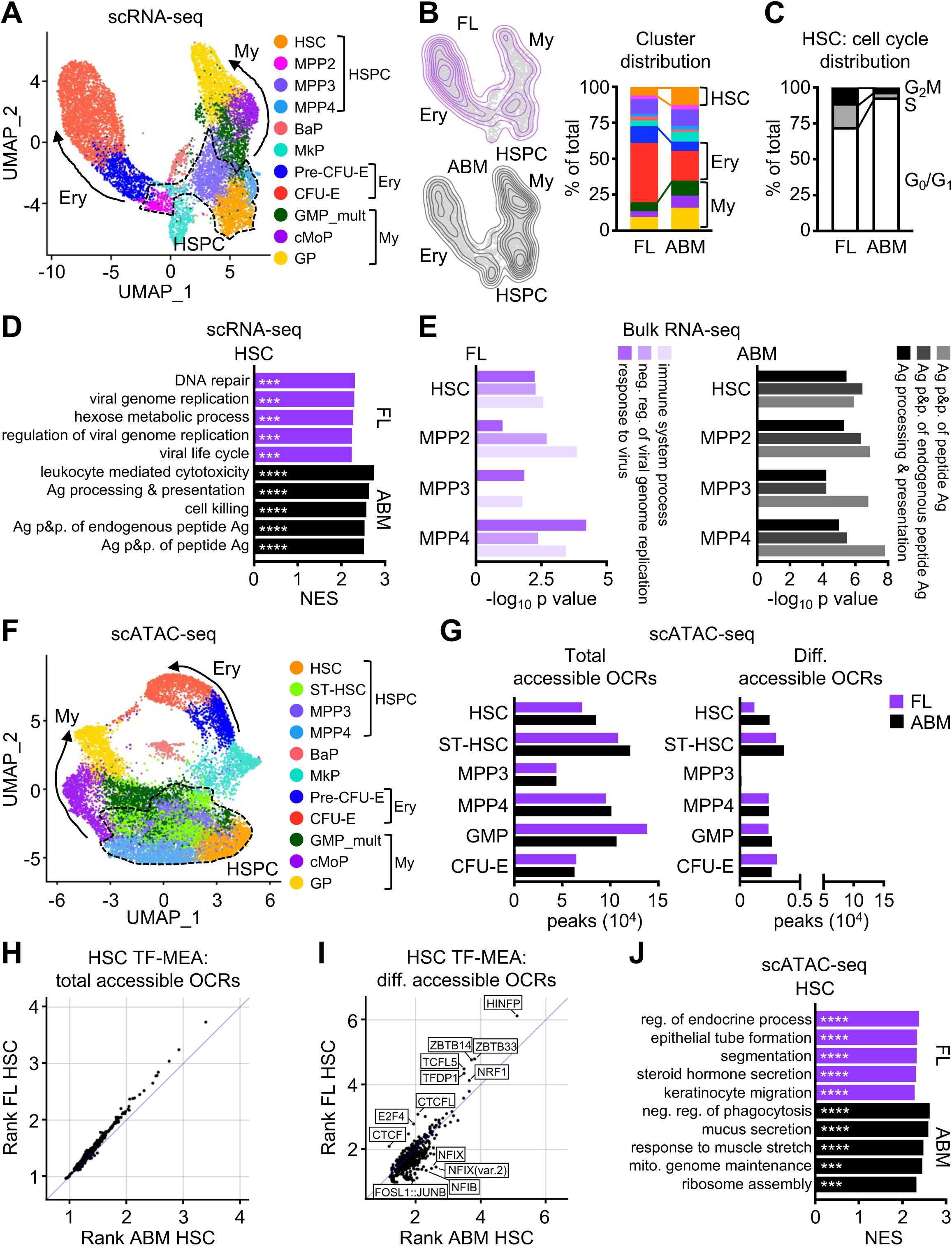
Similar transcriptional landscape in fetal and adult HSPCs. **A)** scRNA-seq UMAP of integrated FL and ABM Lin^-^/c-Kit^+^ (LK) cells with clusters identified by marker genes. Results are from a set of pooled fetuses and adult mice and include 4,909 FL and 7,304 ABM LK cells. **B)** Density projection of cells on UMAPs (left) and distribution of clusters (right) from FL and ABM scRNA-seq. **C)** Cell cycle distribution in HSC cluster from FL and ABM scRNA-seq. **D)** Top 5 pathways differentially enriched in HSC cluster from FL and ABM scRNA-seq (GSEA on DEGs; log_2_ FC > 1, FDR < 0.05); Ag, antigen; p&p., processing and presentation. **E)** Top 3 pathways differentially enriched in FL (left) vs. ABM (right) HSPC bulk RNA-seq (GO/DAVID analyses on DEGs; log_2_ FC > 1, FDR < 0.05). **F)** scATAC-seq UMAP of integrated FL and ABM LK plus Lin^-^/Sca-1^+^/c-Kit^+^ (LSK) cells with clusters identified by marker genes. Results are from a set of pooled fetuses and adult mice and include 6,704 FL and 8,288 ABM LK+LSK cells. **G)** Number of total (left) and differentially (diff., right) accessible open chromatin regions (OCRs; min.pct 0.05, log_2_FC > 0.25, p_adj_ < 0.05) found in the indicated clusters from FL and ABM scATAC-seq. **H)** Transcription factor motif enrichment analysis (TF-MEA) found in total accessible OCRs in FL and ABM HSC clusters. **I)** TF-MEA found in differentially accessible OCRs in FL and ABM HSC clusters. **J)** Top 5 pathways driven by genes closest to OCRs found differentially enriched in HSC cluster from FL and ABM scATAC-seq (GSEA on differentially accessible ORCs); neg., negative; mito., mitochondrial. ***p_adj_ ≤ 0.001; ****p_adj_ ≤ 0.0001. See also Figure S3.

### Unchanged molecular landscape between fetal and adult HSPCs

To address whether differences in chromatin accessibility may underpin the baseline difference in myeloid output of fetal and adult HSPCs, we performed single cell assay for transposase-accessible chromatin with sequencing (scATAC-seq) on a combination of LK and LSK cells obtained from fetal and adult mice. UMAP and cluster identification of the integrated scATAC-seq dataset revealed a similar organization as the scRNA-seq dataset, with a common HSPC-containing center producing both erythroid progenitor and myeloid progenitor arms (**Figure 3F**). Examination of the total number of accessible peaks in each cluster demonstrated between 40,000 and 140,000 open chromatin regions (OCR), with similar numbers present in both fetal and adult clusters, and fewer than 5,000 differentially accessible peaks in either fetal or adult clusters across all HSPC populations (**Figure 3G**). Transcription factor motif enrichment analysis (TF-MEA) of all OCRs showed no differences between fetal and adult HSCs (**Figure 3H**). In contrast, TF-MEA of OCRs found in differentially accessible regions revealed enrichment of motifs bound by TFs known to regulate cell cycle progression and chromatin architecture (like HINFP^50^ and CTCF^51^) in fetal HSCs, and to regulate myelopoiesis (like AP-1^52^ and NFI^53^) in adult HSCs (**Figure 3I**). However, gene set enrichment analysis (GSEA) driven by genes closest to the differentially accessible OCRs in either fetal or adult HSPCs did not show any significant enrichment for pathways associated with blood lineage commitment (**Figures 3J, S3I**). Altogether, our molecular analyses reveal little difference in the baseline transcriptional programs or chromatin conformation that might encourage adult HSPCs towards or restrict fetal HSPCs from myelopoiesis. This suggests that decreased exposure rather than decreased responsiveness to pro-myeloid factors might be the main driver of restricted fetal myelopoiesis, consistent with our observations that fetal HSPCs can be driven to myelopoiesis as readily as adult HSPCs by inflammatory signals *in vitro*, and that fetal HSCs can adopt the behavior of adult HSCs when inhabiting the adult BM niche upon transplantation *in vivo*.

### Fetal HSPCs do not engage emergency myelopoiesis pathways *in utero*

To explore the notion that the maternal-fetal microenvironment modulates the response of fetal HSPCs to pro-myeloid signals, we first evaluated the ability of fetal HSPCs to engage EM pathways in response to a relevant *in vivo* inflammatory challenge by injecting pregnant dams with lipopolysaccharide (LPS). Depending on the dose and embryonic day of administration, LPS is known to result in either *in utero* demise or preterm delivery^54^. We found that a low dose of LPS (100 μg/kg) injected intravenously at E17.5 resulted in preterm delivery in ∼50% of pregnant mice by 24 hours, and thus chose a 16-hour timepoint to reliably compare PBS and LPS-exposed fetuses. This dose of LPS in adult mice yielded the expected inflammatory response as assessed by CBC analyses, with leukocytosis driven by recruitment of neutrophils to the periphery and Sca-1 upregulation on BM HSPCs (**Figures 4A, S4A**). While fetuses clearly responded to maternal LPS exposure with an increase in white blood cell (WBC) counts by CBC (which we subsequently identified as nucleated RBCs, **see STAR Methods**) and elevated Ter119^+^ erythrocyte levels, we observed no increase in circulating neutrophils as in adult blood (**Figures 4A, S4B, S4C, S4D**). We also did not find Sca-1 to be upregulated on FL HSPCs (**Figures S4A**). These data suggest that maternal LPS administration is sensed by the fetal compartment and mobilizes nucleated RBCs into the periphery, a known response to inflammation also described in adults^55^, but fails to elicit neutrophilia as in adults.

**Figure 4.**
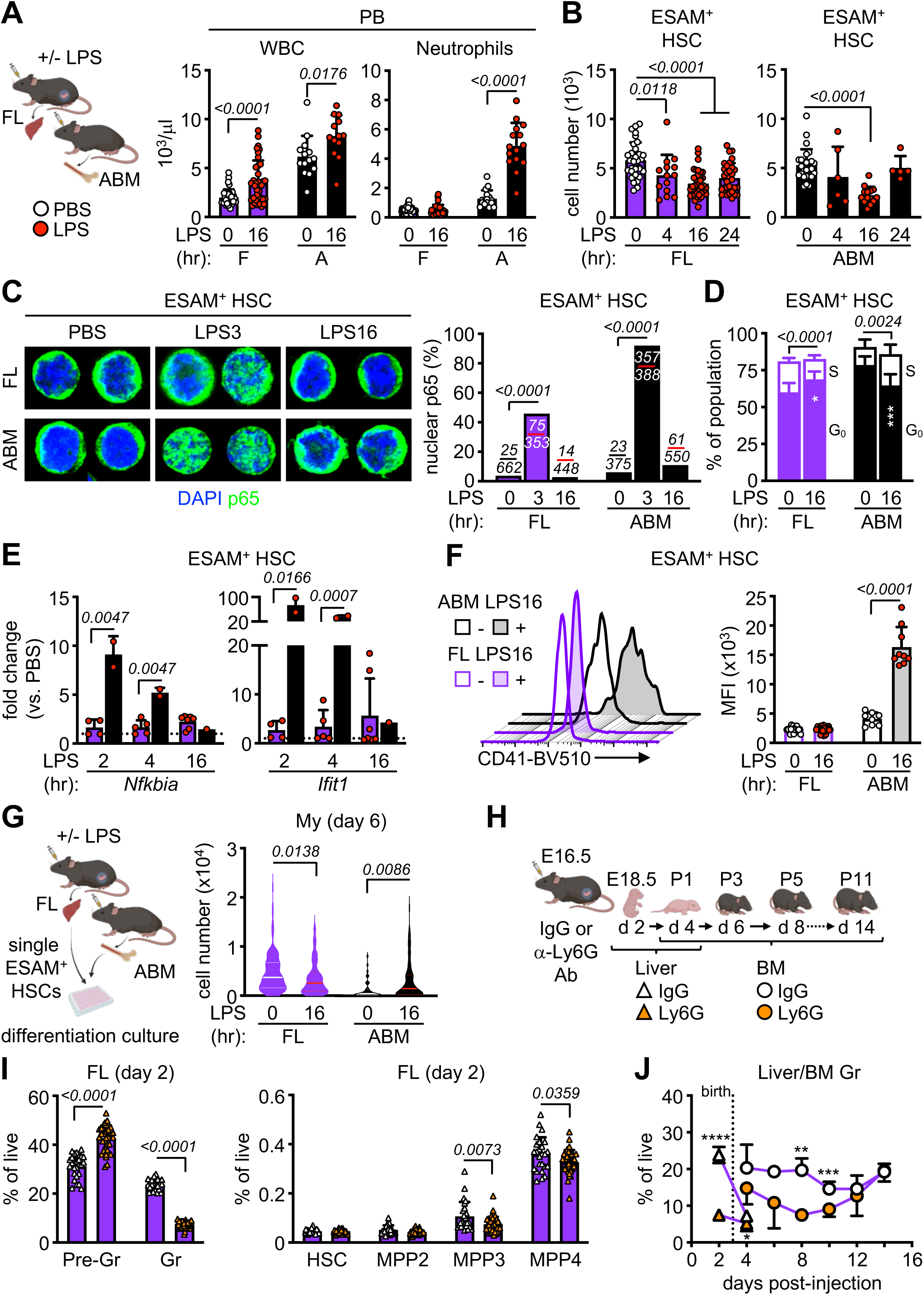
Fetal HSPCs do not engage in emergency myelopoiesis *in vivo*. **A)** Schematic of *in vivo* LPS (100 µg/kg/mouse) experiments with white blood cell (WBC) (left) and neutrophil (right) counts from CBC of PB from LPS-exposed fetal (F) and adult (A) mice (5 independent experiments); hr, hour. **B)** Quantification of LPS-exposed FL (right) and ABM (left) ESAM^+^ HSCs (5 independent experiments). **C)** Representative images (left) and quantification (right) of nuclear p65 in LPS-exposed FL and ABM ESAM^+^ HSCs. Number of cells scored with nuclear p65 vs. all cells counted are indicated over the corresponding bar graph (3 independent experiments). **D)** G_0_ and S phase distribution in LPS-exposed FL and ABM ESAM^+^ HSCs (2 independent experiments in *Fucci2* cell cycle reporter mice). **E)** qRT-PCR expression of *Nfkbia* and *Ifit1* in LPS-exposed FL and ABM ESAM^+^ HSCs (4 independent experiments). **F)** Representative FACS histograms (left) and quantification (right) of CD41 MFI on LPS-exposed ESAM^+^ HSCs (5 independent experiments). **G)** Schematic of single cell differentiation culture of LPS-exposed FL and ABM ESAM^+^ HSCs with cellularity of myeloid (My) colonies at day 6 (3 independent experiments with a range of 37-140 colonies counted per population). Data are shown as violin plots with median and quartiles. **H)** Schematic of IgG or anti-Ly6G antibody injection (0.1 mg/mouse) with time-course of analysis in fetal/neonatal mice. **I)** Quantification of myeloid cells (left) and HSPCs (right) in FL 2 days after IgG or anti-Ly6G exposure (at least 2 independent experiments); Pre-Gr, pre-granulocyte (Mac-1^+^/Gr-1^int^); Gr, granulocyte (Mac-1^+^/Gr-1^+^). **J)** Quantification of granulocytes in the liver (triangles) and BM (circles) of fetal/neonatal mice exposed to IgG or anti-Ly6G during development (at least 2 independent experiments; see Table S1). Dotted line indicates birth. Data are means ± S.D. except for (G); **p_adj_ ≤ 0.01; ***p_adj_ ≤ 0.001; ****p_adj_ ≤ 0.0001. Number of biological replicates and treated fetuses/mice are indicated either in parentheses or with individual circles. See also Figure S4 and Table S1.

We next investigated the HSPC compartment, using the ESAM surface marker to reliably identify LPS-exposed adult and fetal HSCs as ESAM^+^/Flk2^-^/CD150^+^/CD48^-^ LSK (ESAM^+^ HSC) cells^56^ (**Figure S4E**). We observed that LPS stimulation caused a transient drop in adult HSC numbers at 16 hours, followed by recovery by 24 hours, while fetal HSCs were depleted as early as 4 hours post-exposure, with no recovery by 24 hours (**Figure 4B**). We also found a rapid and profound depletion of all fetal MPPs upon LPS exposure (**Figure S4F**). Moreover, LPS-exposed fetal and adult HSCs both exhibited increased NFκB subunit p65 translocation to the nucleus, although the response of fetal HSC was more attenuated (**Figure 4C**). In addition, both HSC populations upregulated PU.1 activity to similar extent (**Figure S4G**), indicating that the ability of fetal HSCs to sense LPS was intact but did not lead to engagement of EM pathways like in adult HSCs. For instance, while LPS induced adult HSCs to enter the cell cycle, increasing the fraction of cells in S phase, it had the opposite effect on fetal HSCs, blocking proliferation and increasing the fraction of cells in quiescent G_0_ phase (**Figure 4D**). LPS also caused upregulation of NFκB (*Nfkbia*) and interferon (*Ifit1*) pathway genes in adult but not fetal HSCs (**Figure 4E**), and increased expression of the myeloid-biased HSC marker CD41 on adult but not fetal HSCs (**Figure 4F**). Finally, while LPS increased the cellularity of myeloid-derived colonies obtained from adult HSCs in single cell *in vitro* differentiation assay, it resulted in the opposite effect on fetal HSCs, with significantly decreased myeloid cellularity (**Figure 4G**). These results demonstrate that while fetuses can sense maternally derived inflammation, and that the initial response of fetal HSCs is intact, they appear restricted from translating those signals into enhanced myelopoiesis.

To overcome the high preterm delivery rate associated with *in utero* LPS exposure that precludes following the fetal HSPC response over time, we employed another method to engage EM pathways and injected pregnant dams with anti-Ly6G antibody to deplete neutrophils and trigger demand-adapted myelopoiesis^35–37^ (**Figure 4H**). In adult mice, granulocyte depletion occurred rapidly by day 2 in the BM and triggered an expansion of all MPP populations followed by a subsequent burst of mature neutrophil production that peaked at day 10 in the PB (**Figures S4H, S4I**). In pregnant dams, anti-Ly6G antibody injected at E16.5 was able to cross the placenta, leading to a robust depletion of granulocytes in the FL 2 days later at E18.5 (**Figure 4I**). However, fetal granulocyte depletion resulted instead in a contraction of MPP3 and MPP4 populations and no outburst of mature neutrophils in the fetal PB, with a slow recovery of myeloid homeostasis over the following 2 weeks after birth in neonatal PB (**Figures 4I**, **4J**, **S4J**). Collectively, these findings show that fetal HSPCs sense EM-initiating insults but fail to activate EM pathways, demonstrating that fetal restriction of myelopoiesis extends to conditions of increased demand like inflammation.

### Molecular analysis confirms lack of EM pathway activation from fetal HSPCs

To investigate the molecular response to EM engagement, we performed scRNA-seq on LK cells isolated from fetal and adult mice exposed to PBS or LPS for 3 or 16 hours. UMAP representation of the integrated results and density projection in each individual dataset confirmed more significant depletion of HSPCs in LPS-treated fetuses compared to adults at both 3 and 16 hours (**Figures 5A, 5B**), with increased molecular signature of quiescence in LPS-exposed fetal HSCs compared to cell cycle entry in LPS-exposed adult HSCs (**Figure S5A**). Comparison of the numbers of DEGs showed a significant transcriptional response in adult HSPCs that was attenuated in fetal HSPCs (**Figure S5B**). At 3 hours, GSEA of LPS-exposed fetal HSPCs revealed enrichment for pathways like nucleosome organization that did not reflect any lineage priming, while GSEA of LPS-exposed adult HSPCs identified pathways such as NFκB signaling consistent with an early response to inflammation (**Figures S5C, S5D**). At 16 hours, we observed an overlap in many negatively enriched pathways like aerobic respiration and cytoplasmic translation, as well as downregulation of genes like *Uqcrh*, a component of the mitochondrial electron transport chain, in both LPS-exposed fetal and adult HSPCs (**Figures 5C, S5D**), suggesting some degree of shared transcriptional repression. However, positively enriched pathways were divergent, with pathways upregulated in LPS-exposed fetal HSPCs including those related to intracellular signaling and cell migration, and in LPS-exposed adult HSPCs those involved in inflammatory response and myelopoiesis driven in large part by upregulation of genes like *Bst2*, which is required for inflammation-mediated HSC activation^57^ (**Figure 5C, S5C, S5E**). LPS-exposed fetal HSCs also showed delayed upregulation of *Spi1* mRNA compared to adult HSCs and lacked activation of PU.1 downstream target genes like *Csf3r* or *Junb* (**Figure S5F**). In addition, LPS-exposed fetal HSCs had repressed *Myc* levels in contrast to adult HSCs that upregulated *Myc* in response to LPS (**Figure S5E**), which we further confirmed using LPS-treated *Myc-eGFP* reporter mice (**Figure S5G**). As a consequence, following LPS exposure, we observed significant enrichment of the Hallmark “*Myc_v1 targets*” module score in the myeloid progenitor wing of the adult UMAP, which was considerably attenuated in the myeloid arm of the fetal UMAP (**Figure 5D**). Together, these results demonstrate that fetal HSPCs respond to maternal inflammation with a transcriptional response that does not include activation of EM pathways as seen in adult HSPCs.

**Figure 5.**
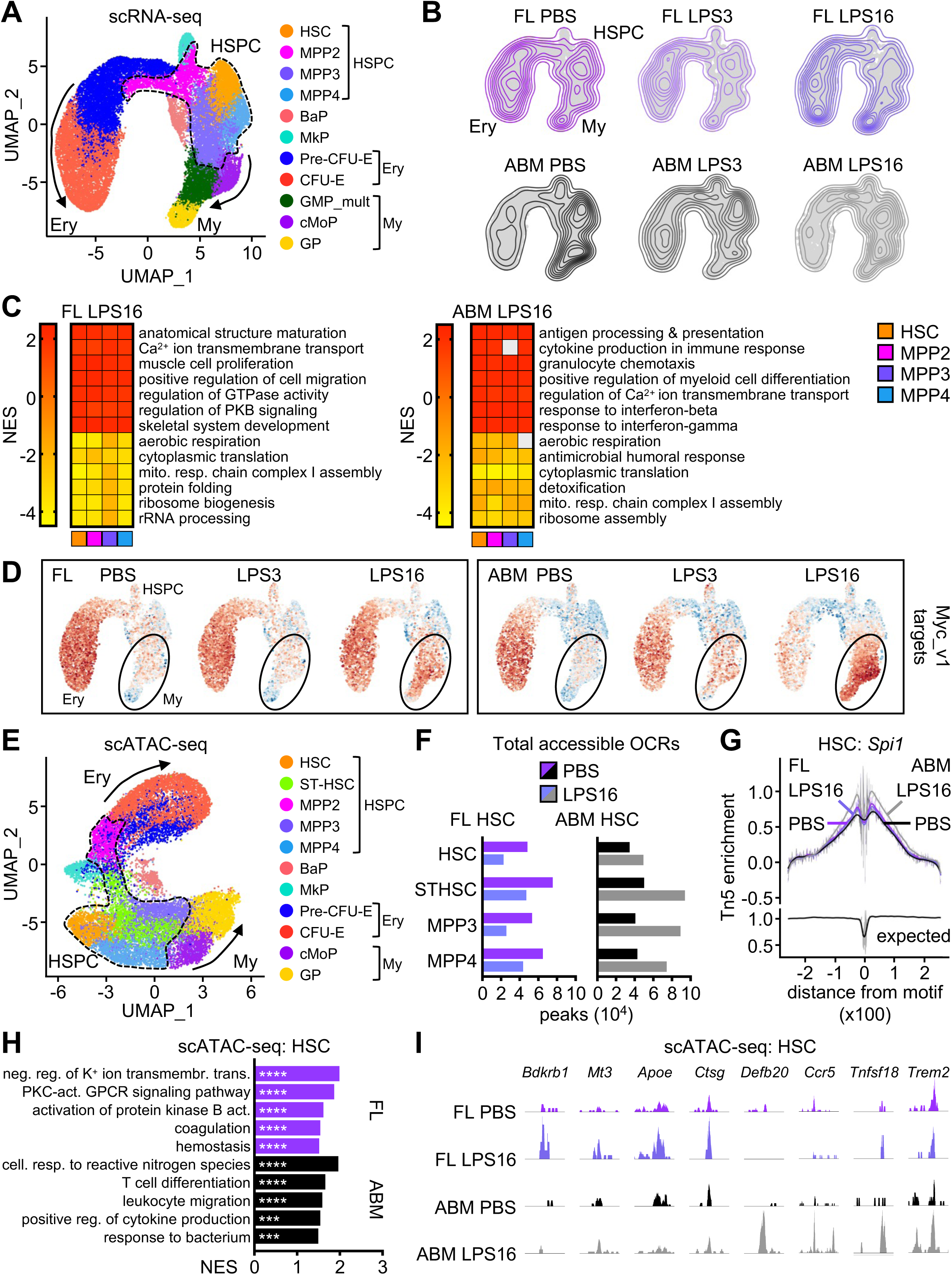
Molecular analysis confirms lack of EM pathway activation from fetal HSPCs. **A)** scRNA-seq UMAP of integrated FL and ABM Lin^-^/c-Kit^+^ (LK) cells isolated from mice exposed to PBS or LPS for 3 (LPS3) or 16 (LPS16) hours with clusters identified by marker genes. Results are from a set of pooled fetuses and adult mice and include 13,995 FL (PBS: 4,947; LPS3: 5,313; LPS16: 3,735) and 15,671 ABM (PBS: 7,181; LPS3: 4,644; LPS16: 3,846) LK cells. **B)** Density projection of cells on UMAPs from LPS-exposed FL and ABM scRNA-seq. **C)** Top pathways differentially enriched in FL (left) and ABM (right) HSPC clusters from 16 hours LPS-exposed FL and ABM scRNA-seq (GSEA on DEGs; log_2_ FC ± 0.25, min.pct 0.25). Grey boxes are pathways not meeting the p_adj_ < 0.05 cut-off; mito. resp., mitochondrial respiratory. **D)** Projection of module score derived from Hallmark “Myc_v1 targets” pathway onto LPS-exposed FL and ABM scRNA-seq UMAPs. **E)** scATAC-seq UMAP of integrated FL and ABM LK cells isolated from mice exposed to PBS or LPS for 16 hours with clusters identified by marker genes. Results are from a set of pooled fetuses and adult mice and include 6,404 FL (PBS: 4,618; LPS16: 1,786) and 11,261 ABM (PBS: 4,768; LPS16: 6,493) LK cells. **F)** Total open chromatin regions (OCRs; min.pct 0.05, log_2_FC > 0.25, p_adj_ < 0.05) in HSC cluster of 16 hours LPS-exposed FL (left) and ABM (right) scATAC-seq. **G)** Transcription factor footprint of *Spi*1 in HSC cluster of 16 hours LPS-exposed FL and ABM scATAC-seq. **H)** Top 5 pathways driven by genes closest to OCRs found differentially enriched in HSC cluster from 16 hours LPS-exposed FL and ABM scATAC-seq (GSEA on differentially accessible ORCs; log_2_ FC ± 0.25, min.pct 0.25); neg., negative; reg., regulation; transmembr., transmembrane; trans., transport; act., activity; cell., cellular; resp., response. I) Examples of scATAC-seq Integrative Genomics Viewer (IGV) tracks in FL and ABM HSC cluster for genes associated with gained OCRs in response to 16 hours LPS exposure. ***p_adj_ ≤ 0.001; ****p_adj_ ≤ 0.0001. See also Figure S5.

To determine whether changes in chromatin accessibility also contribute to the inability of fetal HSPCs to engage EM pathways in response to inflammation, we performed scATAC-seq on LK cells isolated from fetal and adult mice exposed to PBS or LPS for 16 hours (**Figure 5E**). We observed an increase in accessibility and OCRs in LPS-exposed adult HSPCs in contrast to an overall decrease in accessibility and OCRs in LPS-exposed fetal HSPCs (**Figure 5F**). Accordingly, TF-MEA showed enhanced accessibility of *Spi1* motifs in adult but not fetal HSCs (**Figure 5G**), congruent with the divergent PU.1 target gene expression at each developmental stage upon LPS exposure. GSEA showed enrichment for cytokine production and response to bacterium pathways in LPS-exposed adult HSCs, which were driven by increased accessibility at loci such as *Defb20*, *Ccr5*, *Tnfsf18*, and *Trem2*, while pathways enriched in LPS-exposed fetal HSCs involved GPCR and protein kinase B signaling, driven by increased accessibility at loci like *Bdkrb1* and *Mt3*, as well as coagulation and hemostasis, driven by increased accessibility at loci such as *Apoe* and *Ctsg* (**Figures 5H, 5I**). Collectively, these analyses demonstrate that while fetal HSPCs do engage in a molecular response to LPS, with changes in transcriptional signaling response and chromatin accessibility consistent with their limited response to inflammation and RBC mobilization, they fail to activate the classical inflammatory-mediated myelopoiesis pathways as adult HSPCs do, thus providing a molecular basis for their failure to mount an EM response *in utero*.

### External factors control the fetal response to maternal inflammation

To identify the mediator of EM repression in the fetal environment, we used a bead-based multiplex assay to interrogate the inflammatory milieu elicited in response to LPS in both maternal and fetal serum (**Figure 6A**). As expected, we observed strong induction of many cytokines and chemokines in maternal serum in response to LPS (**Figures 6B**, **S6A, S6B**), with the strongest induction occurring by 4 hours (**Figure 6C**). In contrast, none of the same factors were significantly increased in fetal serum in response to LPS, with some other cytokines like IFNβ and CXCL13 being actually decreased (**Figures 6B**, **S6A, S6B**). Consistent with the dispensability of direct cytokine signaling in the fetus, genetic ablation of either IL-1 receptor in *Il1r1^-/-^* fetuses or type I IFN receptor in *Ifnar^-/-^* fetuses had no discernible impact on the fetal response to maternal LPS exposure, with equivalent HSC depletion in the FL and nucleated RBC elevation in the PB of receptor-deleted or-sufficient fetuses (**Figures S6C**, **S6D**). In contrast, we found that the maternal ability to sense inflammatory signaling controlled most of the fetal HSPC response to inflammation. In fact, *Ifnar^-/-^* fetuses exposed to LPS carried by dams that were also *Ifnar^-/-^* had no depletion of HSCs or downstream MPP2 or MPP4 anymore (**Figure 6D**). To further explore this maternal-fetus crosstalk, we focused on TNFα, a known myelopoiesis stimulator^33^ that is significantly upregulated in maternal serum in response to LPS (**Figure 6B**). We set up mixed genotype breeding pairs to generate mother-fetus dyads with TNFα-deficient fetuses carried by TNFα-sufficient mothers and vice versa. Consistently, we found that fetal TNFα was dispensable for the HSC depletion and nucleated RBC enrichment seen in LPS-exposed fetuses, and that maternal TNFα was a necessary driver for those fetal responses to LPS (**Figures 6E, 6F, S6E**). These results reinforce the notion that fetal HSPCs are responsive to external cues and suggest a key role for maternal-derived extrinsic factors in regulating the response of the fetal HSPC compartment.

**Figure 6.**
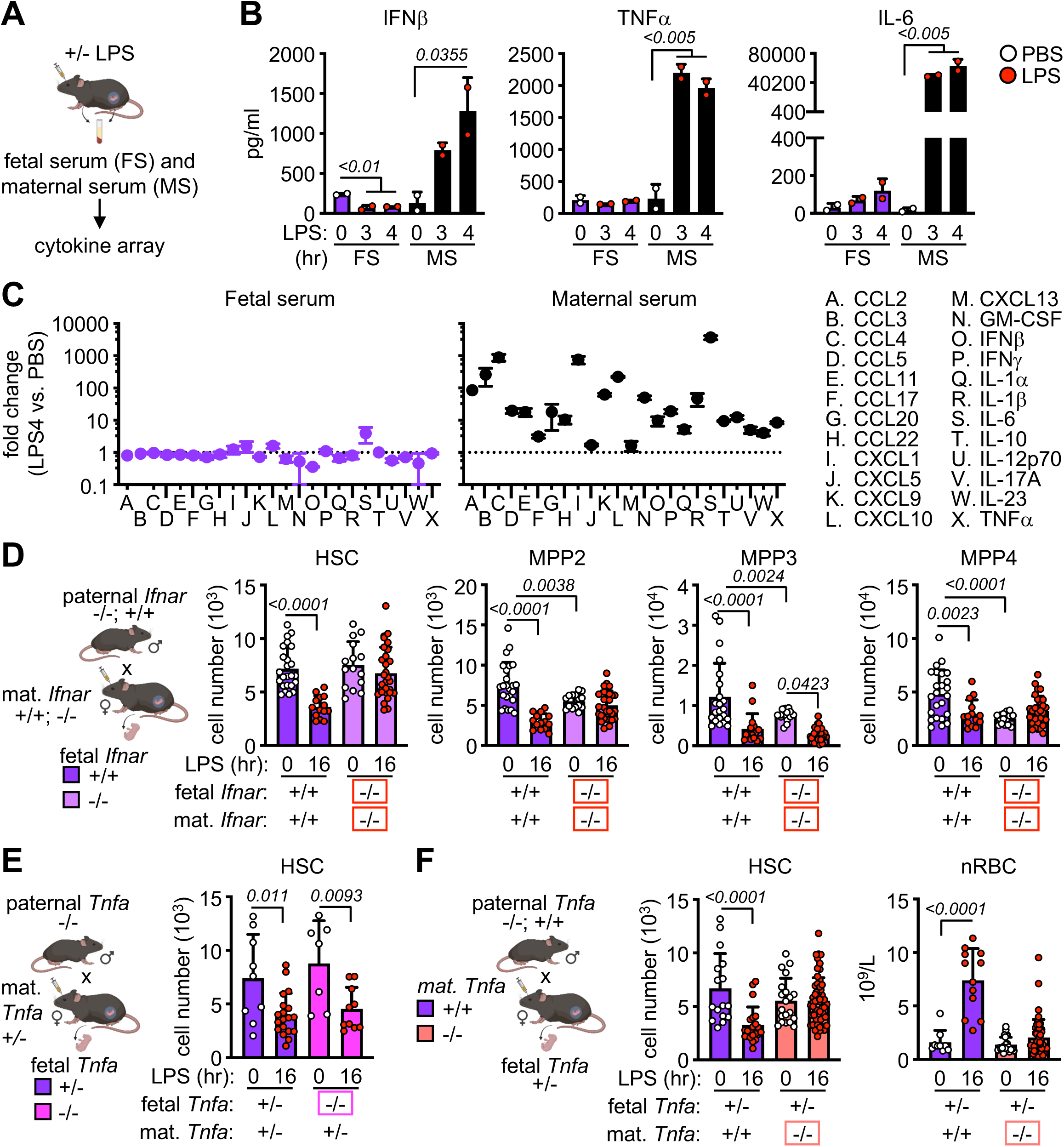
External factors control fetal response to maternal inflammation. **A)** Schematic of *in vivo* LPS exposure and collection of fetal serum (FS) and maternal serum (MS) in pregnant mice. **B)** Cytokine levels in paired fetal and maternal serum from mice exposed to LPS for 3 hours and 4 hours. Circles represent the mean of 2 technical replicates (2 independent experiments). Fetal serum is pooled from all fetuses in the litter (average 5 to 8 fetuses per litter). **C)** Fold change in cytokine/chemokine level in paired fetal (left) and maternal (right) serum from mice exposed to LPS for 4 hours (2 independent experiments). **D)** Schematic of *Ifnar* time-mated pregnancies with quantification of HSPCs in LPS-exposed wild-type and *Ifnar*^-/-^ FL carried by wild-type and *Ifnar*^-/-^ dams (3 independent experiments); mat., maternal. **E)** Schematic of *Tnfa* time-mated pregnancies with quantification of HSCs in LPS-exposed *Tnfa*^+/-^ and *Tnfa*^-/-^ FL (4 independent experiments). **F)** Schematic of *Tnfa* time-mated pregnancies with quantification of FL HSCs (left) and PB nucleated RBCs (nRBC, right) in LPS-exposed *Tnfa*^+/-^ fetuses carried by *Tnfa*^+/+^ or *Tnfa*^-/-^ dams (5 independent experiments). Data are means ± S.D. Number of biological replicates and treated fetuses/mice are indicated with individual circles. See also Figure S6.

### Maternal IL-10 restricts fetal EM activation

To directly test whether the predominantly anti-inflammatory milieu of late pregnancy^7^ could have the effect of restricting fetal myelopoiesis, we focused on IL-10, a cytokine known to play a major role in pregnancy tolerance^58^ and that we found upregulated in maternal serum upon LPS exposure (**Figure 6C**). We made use of *Il10^-/-^*mice^59^ to set up mixed maternal-fetal dyads in which IL-10 sufficient (*Il10^+/-^*) fetuses were carried by either IL-10-sufficient *Il10^+/+^* dams (fetus^WT-mom^) or IL10-deficient *Il10^-/-^* dams (fetus^KO-mom^) (**Figure 7A**). While we found no difference in the pregnancy and delivery rate from *Il10*^-/-^ dams in normal conditions, they became visibly sick following LPS treatment with hunched posture, reduced activity, and rapid induction of *in utero* fetal demise upon 8 hours LPS exposure (**Figure S7A**). Focusing on 4 hours post-LPS injection, we observed that in the absence of maternal IL-10 in fetus^KO-mom^, HSCs were not depleted and showed a modest but significant upregulation of the myeloid-biased marker CD41 on their surface, and that all MPP populations were expanded instead of being contracted as in fetuses^WT-mom^ (**Figures 7B, 7C, S7B**). These effects were specific to maternal IL-10 as we found no difference in the response to LPS between *Il10^+/-^* and *Il10^-/-^* fetuses carried by *Il10^+/-^* dams mated to *Il10^-/-^* sires (data not shown). We next used our Ly6G depletion model to follow the kinetics of this response, finding that maternal anti-Ly6G antibody administration did not compromise the pregnancy of *Il10^-/-^* females and depleted fetal granulocytes equally effectively in fetuses^WT-mom^ and fetuses^KO-mom^ (**Figure 7D**). However, in contrast to the lack of EM engagement in fetuses^WT-mom^, we now observed expansion of every EM-involved HSPC in day 2 FL from fetuses^KO-mom^ (**Figure 7E**), and much more robust mature myeloid cell production in the BM and PB of neonates carried by Ly6G-treated *Il10^-/-^*dams than WT dams (**Figures 7F, S7C**). To confirm that the absence of maternal IL-10 altered the inflammatory milieu, we also tested the composition of maternal and fetal serum in LPS-exposed mixed *Il10* genotype dyads (**Figure S7D**). As expected, maternal serum from *Il10^-/-^* dams showed an enhanced inflammatory response to LPS compared to WT dams (**Figure S6A**), which now translated into significant increases in TNFα, IL-6, CXCL1, and CXCL5 in fetal serum (**Figure S7D, S7E**). These results suggest that by dampening inflammation, maternal IL-10 prevents the engagement of EM pathways in fetal HSPCs.

**Figure 7.**
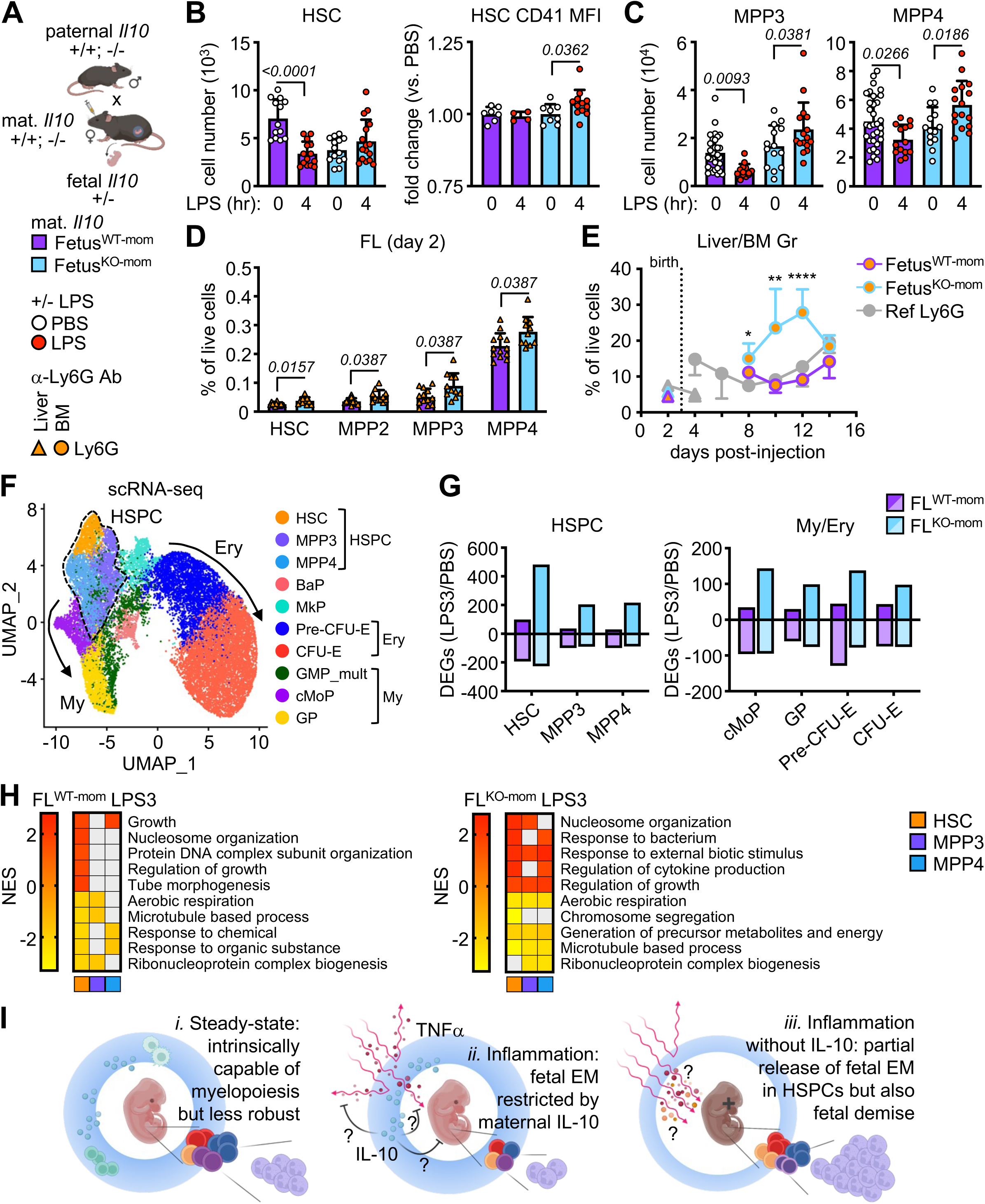
Maternal IL-10 restricts fetal response to inflammation. **A)** Schematic of *Il10* time-mated pregnancies with naming convention and treatments; fetus^WT-mom^, *Il10^+/-^* fetuses carried by wild-type dams; fetus^KO-mom^, *Il10^+/-^* fetuses carried by *Il10^-/-^* dams. **B)** Quantification of FL HSCs (left) and fold change of CD41 MFI on HSCs (right) in fetus^WT-mom^ and fetus^KO-mom^ mice exposed to LPS for 4 hours (6 independent experiments). **C)** Quantification of FL MPP3 (left) and MPP4 (right) in fetus^WT-mom^ and fetus^KO-mom^ mice exposed to LPS for 4 hours (6 independent experiments). **D)** Quantification of FL HSPCs 2 days after anti-Ly6G exposure in fetus^WT-mom^ and fetus^KO-mom^ mice (2 independent experiments). **E)** Quantification of granulocytes in the liver (triangles) and BM (circles) of fetal/neonatal fetus^WT-mom^ and fetus^KO-mom^ mice exposed to anti-Ly6G during development (at least 2 independent experiments; see Table S3). Dotted line indicates birth and grey line the reference production of granulocytes in Ly6G-treated WT dams. **F)** scRNA-seq UMAP of integrated FL LK cells isolated from fetus^WT-mom^ and fetus^KO-mom^ mice exposed to PBS or LPS for 3 hours with clusters identified by marker genes. Results are from a set of pooled fetuses and include 12,594 fetus^WT-mom^ (PBS: 6,209; LPS3: 6,385) and 11,890 fetus^KO-mom^ (PBS: 4,977; LPS3: 6,913) FL LK cells. **G)** Total number of DEGs in HSPC (left) and My/Ery (right) clusters from 3 hours LPS-exposed fetus^WT-mom^ and fetus^KO-mom^ FL scRNA-seq.. **I)** Top pathways differentially expressed in fetus^WT-mom^ and fetus^KO-mom^ HSPC clusters from 3 hours LPS-exposed FL scRNA-seq (GSEA on DEGs; log_2_ fold change +/-0.25, min.pct 0.25). Grey boxes are pathways not meeting the p_adj_ < 0.05 cut-off. **J)** Model of shielding of fetal emergency myelopoiesis by the maternal anti-inflammatory milieu. At steady state (*i.*), fetal HSPCs are intrinsically capable of myelopoiesis but their myeloid cell output is less robust than that of adult HSPCs. In response to inflammation (*ii.*), fetal HSPCs fail to activate EM pathways, restricted by maternal IL-10. In the absence of maternal IL-10 (*iii.*), there is partial restoration of EM in fetal HSPCs at both a cellular and transcriptional level, but at the cost of fetal demise. Data are means ± S.D.; *p ≤ 0.05; **p ≤ 0.01; ****p ≤ 0.0001. Number of biological replicates and treated fetuses/mice are indicated with individual circles. See also Figure S7.

To directly address whether maternal IL-10 suppresses fetal HSPC activation, we performed scRNA-seq on LK cells isolated from fetuses^WT-mom^ and fetuses^KO-mom^ exposed to LPS for 3 hours (**Figure 7G**). Examination of DEGs showed a strikingly enhanced transcriptional response in LPS-exposed fetuses^KO-mom^ compared to fetuses^WT-mom^, with near restoration of the number of upregulated DEGs in the HSC cluster (483) (**Figure 7H**) to that observed in adult HSC cluster (662) after 3 hours LPS exposure (**Figure S5B**). GSEA also revealed upregulation of multiple pathways consistent with an early inflammatory response in HSPC clusters from LPS-exposed fetuses^KO-mom^ (**Figure 7I**). This included restoration of pathways induced by LPS in adult HSPC clusters such as response to bacterium, response to external biotic stimulus, etc., and induction of genes like *Ccl4*, *Pde4b*, or *Nfkbia* that were also shared with LPS-exposed adult HSC clusters (**Figures S7D, S7E**), suggesting a *bona fide* restoration of EM pathways. Collectively, these results demonstrate that loss of maternal IL-10 restores EM activation in fetal HSPCs but at the cost of premature parturition. They confirm that extrinsic maternal factors that have likely evolved to protect pregnancy also function to restrict the fetal response to inflammation, hence limiting the ability of fetal HSPCs to engage EM (**Figure 7J**) and directly contributing to perinatal neutropenia and sepsis.

## DISCUSSION

Here, we provide a comprehensive roadmap of hematopoietic development during the transition from perinatal to adult stages in the mouse. By profiling HSC and MPP populations phenotypically, functionally, and molecularly, we show that while the structure of the early hematopoietic hierarchy is conserved from the late fetal period to adulthood, fetal HSPCs have a baseline deficit in myelopoiesis. Using a clinically relevant model of prenatal inflammation, we demonstrate that fetal HSPCs are restricted from engaging EM by cell-extrinsic factors and identify maternal IL-10 as one such key factor limiting myeloid cell output *in utero*. These results advance our understanding of the vastly understudied period of perinatal hematopoiesis, establishing cell-extrinsic restriction of EM pathway activation as a key driver of neonatal neutropenia.

We establish that fetal HSPCs have limited myeloid cell output relative to adult HSPCs, which is a lineage-specific deficit and not a general defect because fetal erythropoiesis and lymphopoiesis are intact and even enhanced compared to adult output. Our results uncover important distinctions between proliferative capacity, lineage potential, and lineage output between fetal and adult populations. We confirm the well-established observation that fetal HSCs have increased proliferative capacity compared to adult HSCs^40–42^, and, for the most part, extend this observation to fetal MPPs. Most importantly, we establish that while fetal HSPCs are actively restricted in their baseline production of myeloid cells, they do have equivalent myeloid lineage “potential” as adult HSPCs. First, we found no transcriptional poising or chromatin architecture at steady state in myeloid lineage genes that preclude fetal HSPCs from engaging in myelopoiesis. Second, we demonstrate using genetic approaches that the dominant transcriptional signatures distinguishing fetal from adult HSPCs we and others identified^46–48^ (ISGs in the fetus and antigen processing/presentation in the adult) do not influence myeloid cell production. And third, we establish that fetal HSPCs have the capability of producing myeloid cell to the same extent as adult HSPCs in response to strong pro-myeloid stimuli both *in vitro* and *in vivo*. These findings establish that fetal and adult HSPCs have similar potential for myelopoiesis, which does not necessarily translate into equivalent myeloid cell output. In fact, at baseline, we found that fetal HSCs and MPPs produce fewer myeloid cells than their adult counterparts, which occurs without comparable deficits in erythroid or lymphoid cell production and can be read out in both *in vitro* culture and *in vivo* transplantation assays. This diminished production is already apparent at multiple levels of the fetal hematopoietic hierarchy, with fewer myeloid-biased CD41^+^ HSCs, MPP3 and FcγR^+^ MPP3, and GMPs, compared to adult. These results are concordant with the majority of reports showing either an erythroid or lymphoid bias to fetal hematopoiesis^60–62^, although none of these prior studies specifically evaluated the myeloid output of fetal hematopoiesis. In this context, our results demonstrate that despite increased proliferative capacity and equivalent myeloid potential, fetal HSPCs are significantly restricted in their ability to produce myeloid cells compared to their adult counterparts.

We further establish that myelopoiesis in response to inflammation, so-called “emergency myelopoiesis”, is not activated in fetal HSPCs. We use LPS to mimic bacterial infection, which in the adult has been shown to activate HSCs and trigger myelopoiesis^63^, likely through both direct^64,65^ and indirect^66^ mechanisms. However, we find that in response to LPS, fetuses fail to activate myelopoiesis, with no recruitment of neutrophils in the circulation or activation of EM pathways in fetal HSPCs. We speculate that the lack of induction of PU.1 and the resulting lack of assembly of a regulatory network involving AP-1 and Myc activation could be at the heart of the lack of engagement of EM transcriptional response in fetal HSPCs, a mechanism that remains to be fully explored. At E18.5, the hematopoietic compartment is also starting to shift from hepatic to medullary, with HSPCs identifiable in both FL and fetal BM. We focused our analysis of the HSPC compartment to the FL, as the vast majority of HSPCs are still hepatic at this point, a decision validated by the recent observation that fetal BM HSPCs contribute minimally to blood output during embryogenesis and do not acquire functionality until after birth^23^. Furthermore, the absence of LPS-induced neutrophilia in fetal blood suggests that EM pathways are not being activated in fetal HSPCs in either location. One trivial possibility to explain this observation could be fetal “ignorance” of maternal inflammation. However, maternally administered LPS to pregnant mice has been widely used to model maternal immune activation and subsequent neurodevelopmental impairment in offspring^67^, clearly establishing that fetuses can sense LPS-induced maternal inflammation. We also provide evidence demonstrating that fetuses are not ignorant of maternal inflammation, including intact NF-kB p65 nuclear localization in fetal HSCs and both transcriptional activation and chromatin accessibility changes in each fetal HSPC population. Instead, we found that maternal inflammation increases the proportion of quiescent (G_0_) HSCs, uniformly depletes HSCs and MPPs, and elicits transcriptional responses that favor alternative pathways (e.g., coagulation) rather than upregulation of myeloid pathway genes. Employing a second model of demand-adapted myelopoiesis by depleting granulocytes with α-Ly6G antibody, we followed the kinetics of this response across the perinatal period, showing that the lack of EM engagement in fetal HSPCs translated into absence of the burst of mature myeloid cells characteristic of EM activation in the adult. While some of our observations are in contrast to two recent studies demonstrating transient expansion and activation of a subset of fetal HSCs as well as MPP4 leading to persistence of fetal-like lymphoid cells in the neonate^68,69^, it is likely that differences in the types of inflammatory stimuli, embryonic timing of exposure, and route of administration will account for these discrepancies. This emphasizes the need for continued investigations of fetal hematopoiesis *in situ*, and for further integration of the results obtained from different challenges. Our findings show that fetal HSPCs are actively restricted from engaging EM pathways and amplifying myeloid cell production in response to inflammation, an observation that has significant implications for understanding the mechanisms driving sepsis-induced neonatal neutropenia.

We demonstrate that the restriction of emergency myelopoiesis *in vivo* is primarily regulated by cell-extrinsic mechanisms, and identified maternal IL-10 as a key inhibitor of EM pathway activation in fetal HSPCs. IL-10 is a crucial regulator of immune tolerance during pregnancy^58^. It is dispensable during unperturbed pregnancy^70^ but is critical for suppressing the production of pro-inflammatory cytokines, nitric oxide, and prostaglandins in response to low-grade inflammation, which then lead to placental infiltration, myometrial contraction, intrauterine fetal demise, and preterm delivery^71–73^. Multiple cells have been proposed to be the local source of IL-10 during pregnancy and the relevant IL-10-responsive cells^74,75^. Our studies clearly identify maternal, rather than fetal (or placental), IL-10 as crucial for restricting fetal EM, finding that in its absence, fetal HSPCs are activated and expanded, show molecular evidence of EM pathway engagement, and support enhanced myeloid output. The mechanism of this effect remains to be established, in particular to determine whether maternal IL-10 acts directly on fetal HSPC gene regulation or indirectly on other maternal or fetal-derived cells to control their production of factor(s) that will block EM engagement in fetal HSPCs. While the identification of such factor(s) will be the focus of future studies, our results establish that maternal anti-inflammation constrains EM in the fetus, and that dampening this regulation releases the potential of fetal HSPCs to increase myeloid cell production. This suggests that boosting neonatal capacity for myeloid cell production in the context of inflammation or sepsis-induced neonatal neutropenia might best be achieved by targeting such extrinsic restriction. However, the current cost of removing IL-10 is premature parturition, which illustrates the need for further mechanistic studies to identify actionable targets for translational applications.

While it may seem counterintuitive for the fetus to be specifically restricted from myeloid cell production, we believe this is in fact a bystander effect of the dominant immune-suppressive mechanisms that function to maintain pregnancy until term. We speculate that IL-10 may not be unique in this regard, and that removing any other layers of immune regulation that maintain pregnancy could be sufficient to partially release fetal myelopoiesis, a hypothesis that remains to be tested. In the absence of modern medical interventions, premature delivery of a fetus insufficiently developed to survive *ex utero* is an evolutionary dead-end. Infection is the leading cause of preterm delivery, and there is evidence that tonic (non-pathogenic) inflammation contributes to the initiation of parturition^76,77^. Inflammatory pathways must therefore be tightly regulated prior to delivery, resulting in the necessary restriction of what might otherwise be beneficial responses. It therefore remains to be determined whether dampening anti-inflammatory mechanisms during late pregnancy could be a viable therapeutic strategy to boost myeloid cell production without causing parturition. Human validation studies will also be necessary to determine whether similar mechanisms are in place and can be targeted in a different maternal-fetal interface structure and timing of hematopoietic organ colonization. Our discovery that fetal HSPCs do not have a cell-intrinsic deficit in myeloid cell production suggest that it will be possible to develop interventions to specifically improve their output, which opens an entirely new field of investigations for treating neonatal neutropenia.

## ACKNOWLEDGMENTS

We thank O. Olson for advice on many experiments, J. Perez Bruno for technical assistance, Lee Grimes (Cincinnati’s Children) for Fucci2 reporter mice, M. Kissner and team members for excellent operation of the CSCI Flow Cytometry Core facility, Erin Bush and team members for managing the Single Cell Analysis Core, and all members of the Passegué laboratory for critical insights, support and feedback. A.C. was supported by NIH K08DK124657, J.W.S. by long-term EMBO ALTF-2021-196 and Damon Runyon Cancer Research Foundation (William Raveis Charitable Fund Fellow) DRG-2493-23, M.A.P. by NIH TL1DK136048, C.A.M. by NIH F31HL160207, M.K. by ASH RTAF Award, P.V.D. by NIH F31 HL151140, and E.P. by LLS Scholar Award and NIH R35HL135763. This work was funded by a Louis V. Gerstner Jr. Scholar Award to A.C. and NIH R35HL135763 to E.P., and supported in part through the NIH Cancer Center Support Grant P30CA013696 and NIH UL1TR001873 to the Genomics and High Throughput Screening Shared Resource. The content is solely the responsibility of the authors and does not necessarily represent the official views of the NIH and other funders.

## AUTHOR CONTRIBUTIONS

Conceptualization, A.C. and E.P.; Investigation, A.C., M.A.P., C.A.M., M.K., and C.M.P.; Formal Analysis, J.W.S. and P.V.D.; Writing – Original Draft, A.C. and E.P.; Writing – Review & Editing, A.C., J.W.S., C.A.M., M.K., and E.P.; Funding Acquisition, A.C. and E.P; Supervision, A.C. and E.P.; Project administration, E.P.

## INCLUSION AND DIVERSITY STATEMENT

We support inclusive, diverse, and equitable conduct of research.

## STAR METHODS

### Resource Availability

#### Lead Contact and Materials Availability

Further information and requests for resources and reagents should be directed to and will be fulfilled by the Lead Contact, Emmanuelle Passegué, or co-corresponding author, Amelie Collins.

#### Materials Availability

This study did not generate new unique reagents.

#### Data and code availability

Bulk RNA sequencing, single cell RNA sequencing, and single cell ATAC sequencing have been deposited at GEO and are publicly available as of the date of publication. Accession numbers are listed in the key resources table. This paper does not report original code. Any additional information required to reanalyze the data reported in this paper is available from the lead contact or co-corresponding author upon request.

### Experimental Model and Subject Details

#### Animals

All mice were bred and maintained in mouse facilities at CUIMC in accordance with IACUC protocols approved at the institution. Mice were maintained on a 12 hours light cycle in SPF conditions. B6 (C57BL/6J), BoyJ (B6.SJL-*Ptprc^a^Pepc^b^*/BoyJ), *Ifnar*^-/-^ (B6.129S2-*Ifnar1*^tm1Agt^/Mmjax), *Il1r1*^-/-^ (B6.129S7-*Il1r1*^tm1Imx^/J), *Il10*^-/-^ (B6.129P2-*Il10*^tm1Cgn^/J), *B2m*^-/-^ (B6.129P2-*B2m*^tm1Unc^/DcrJ), *H2*^-/-^ (B6.129S2-*H2*^dlAb1-Ea^/J), and *Myc-eGFP* (Myc^tm1.1Dlev^/J) mice were purchased from the Jackson Laboratory. *β-actin-GFP* (*b-actin* C57Bl/6-Thy1.1) mice^78^ and *PU.1-eYFP* mice^80^ were breed in-house, and *Fucci2* (R26Fucci2aR) reporter mice^79^ were obtained from Lee Grimes (Cincinnati Children’s). Wherever possible, littermate controls were used for genetically modified mice, otherwise age- and sex-matched B6 mice were used as wild-type controls. Embryos were staged as days post coitum, with embryonic day E0.5 considered as noon of the day a vaginal plug was detected after overnight mating. Age of fetal and neonatal mice is indicated throughout the paper. Adult mice were 6-12 weeks unless otherwise indicated. No specific randomization or blinding protocol was used. For a number of experiments, we confirmed the sex of E18.5 fetal by PCR^81^ and observed that female and male fetuses had equivalent HSPC depletion in response to maternal LPS compared to PBS. Thereafter, results from male and female fetuses were pooled, and both male and female animals were used indifferently in the study.

#### Cell lines

OP9 cells^84^ were purchased from ATCC (CRL 2749). OP9 cells were grown on 0.1% gelatin-coated plates in α-Minimal Essential Media (α-MEM, made from powder, Gibco 12000-022) containing 20% FBS (Corning, 35-011-CV) in 5% CO_2_ at 37° and passaged prior to reaching confluence.

### Method Details

#### In vivo assays

For LPS-induced inflammatory challenge, mice were injected once by retro-orbital injection with 0.1 mg/kg lipopolysaccharide (LPS from Escherichia coli O55:B5; Sigma-Aldrich) in 100 µl phosphate-buffered saline (PBS) or PBS control. For granulocyte depletion, mice were injected once by retro-orbital injection with 0.1 mg of anti-Ly6G antibody (BioXcell, BE0075-1) or IgG2a isotype control (BioXCell, BE0089) in 100 µl PBS. For short-term *in vivo* lineage tracing assays, BoyJ recipient mice were sub-lethally irradiated (8.5 Gy, delivered in split doses 3 hours apart) using an X-ray irradiator (Faxitron) and injected retro-orbitally with 2000 donor cells (from B6, *β-actin-GFP*, *Ifnar*^-/-^, *B2m*^-/-^, or *H2*^-/-^ mice) within the next 6 hours. Recipient mice receiving *B2m*^-/-^ (and wild-type control) HSCs were depleted of natural killer cells with 0.1 mg anti-NK1.1 antibody weekly, starting on the day of transplantation. For *in vivo* IL-1β treatment following transplantation, recipient mice were injected intraperitoneally with 0.5 μg IL-1β (Peprotech, 211-11B) in 100 ml PBS/0.2% BSA or PBS/0.2% BSA alone once daily starting on the day of transplantation, for 30 days. For long-term *in vivo* HSC transplantation assays, BoyJ recipient mice were lethally irradiated (10.5 Gy, delivered in split doses 3 hours apart) and retro-orbitally injected with 250 donor HSCs together with 600,000 Sca-1-depleted BoyJ BM cells within the next 6 hours. Irradiation recipient mice were administered polymyxin and neomycin-containing water for 4 weeks following transplantation to prevent opportunistic infection, and chimerism was analyzed over time by repeated bleedings. For chimerism analyses, peripheral blood (PB) was obtained by micro-capillary tube from the retro-orbital plexus and collected in tubes containing 4 ml of 10 mM EDTA-containing ACK (150_jmM NH_4_Cl/10_jmM KHCO_3_) lysis buffer before being washed, lysed again in ACK buffer, washed, and resuspended in primary antibodies. For complete blood count (CBC) analyses, PB was aspirated with a 31G needle from cardiac puncture in adults and neonates or cervical decapitation in fetuses following euthanasia, and transferred to an EDTA-containing tube (Becton-Dickinson) for analyses on a Genesis (Oxford Science) hematology system. We noted that our hemocytometer was not able to differentiate between lymphocytes and nucleated RBCs (nRBC) in fetal blood, a known confounder of leukocyte count by automated hemocytometers^85^. We confirmed that elevated “lymphocytes” in fetal blood were actually nRBCs by both flow cytometry and manual inspection of blood smears. For PB smears, blood was streaked on a Diamond White Glass microscope slide (Globe Scientific, 1380-20) and stained with Wright-Giemsa solution (MilliporeSigma, 742-45). Slides were imaged on an Olympus BX43 microscope with an Olympus DP27 camera, and images were acquired using cellSens Entry imaging software.

#### Flow cytometry

Adult BM cells were obtained by crushing leg, arm, and pelvic bones in staining media composed of Hanks’ buffered saline solution (HBSS) containing 2% heat-inactivated FBS (Corning, 35-011-CV). Fat and tissue of neonatal bones were removed carefully under a dissecting scope before crushing in staining media. For individual fetal liver (FL) or neonatal liver, cells were obtained by mechanical dissociation with a P1000 µl tip. For pooled FL and neonate livers, cells were obtained from tissue initially dissociated between the frosted ends of two microscope slides followed by serially passaging through 18G, 20G, and 21G needles. Spleens were mechanically dissociated between the frosted ends of two microscope slides in staining media. For all samples, RBCs were removed by ACK lysis. For adult/neonatal BM cells, single-cell suspensions of cells were purified on a Ficoll gradient (Histopaque 1119, Sigma-Aldrich). Cellularity was determined with a ViCELL-XR automated cell counter (Beckman-Coulter). All staining was performed on ice for 45 minutes unless otherwise indicated, and cells were blocked with rat IgG (Sigma-Aldrich) for 15 minutes prior to primary stain. For HSPC isolation, cells were pre-enriched for c-Kit^+^ cells using c-Kit microbeads (130-091-224; Miltenyi Biotech) and an AutoMACS cell separator (Miltenyi Biotech). The same lineage cocktail was used for all HSPC analyses and contained B220-PE-Cy5, CD3-PE-Cy5, CD4-PE-Cy5, CD5-PE-Cy5, CD8a-PE-Cy5, Gr-1-PE-Cy5, Mac-1-PE-Cy5, and Ter119-PE-Cy5 for staining of adult and neonatal cells, with CD4-PE-Cy5 and Mac-1-PE-Cy5 excluded for staining of FL cells. For LSK and LK isolation, c-Kit-enriched cells were stained with lineage cocktail, c-Kit-APC-Cy7, and Sca-1-BV421. For HSPC isolation, c-Kit-enriched cells were stained with lineage cocktail, c-Kit-APC-Cy7, Sca-1-BV421, Flk2-PE, CD150-BV650, CD48-AF700, and ESAM-APC. For HSPC quantification, cells were stained with lineage cocktail, c-Kit-APC-Cy7, Sca-1-BV421, Flk2-PE, CD150-BV650, CD48-AF700, CD34-FITC, FcγR-PE-Cy7, ESAM-APC, and CD41-BV510. For HSPC quantification in Fucci2 mice, cells were stained with lineage cocktail, c-Kit-APC-Cy7, Sca-1-BV421, Flk2-bio/SA-BV711, CD150-BV650, CD48-AF700, and ESAM-APC. For HSPC analyses in *PU.1-eYFP* and *Myc-eGFP* mice, cells were stained with lineage cocktail, c-Kit-APC-Cy7, Sca-1-BV421, Flk2-PE, CD150-BV650, CD48-AF700, and ESAM-APC. For MHCI and MHCII expression on HSPCs, cells were stained with lineage cocktail, c-Kit-APC-Cy7, Sca-1-BV421, Flk2-PE, CD150-BV650, CD48-AF700, H2-K^b^-FITC, and I-A/I-E-APC. For mature cell quantification including in PB, cells were stained with B220-APC-Cy7, CD3e-APC, CD19-BV650, Gr-1-BV421, Mac-1-BV786, and Ter119-PE-Cy5 or Ter119-FITC. For PB chimerism analyses from *β-actin-GFP* donor mice, cells were stained with B220-APC-Cy7, CD3e-APC, Gr-1-eF450, Mac-1-PE-Cy7, and Ter119-PE-Cy5. For PB chimerism analyses from all other donors, cells were stained with B220-APC-Cy7, CD3e-APC, CD45.1-PE, CD45.2-FITC, Gr-1-BV421, Mac-1-BV786, and Ter119-PE-Cy5. For single cell differentiation culture, cells were stained with CD45-APC-Cy7, CD71-BUV395, CD41-BV510, CD61-PE, Gr-1-BV421, and Mac-1-BV786. For single cell OP9 differentiation culture, cells were stained with CD71-BUV395, Gr-1-BV421, CD19-BV650, Mac-1-BV786, CD41-FITC, CD61-PE, and NK1.1-PE-Cy5. For *in vitro* myeloid differentiation bulk culture, cells were stained with Sca-1-BV421, c-Kit-APC-Cy7, FcγR-PE-Cy5.5, and Mac-1-PE-Cy7. For *in vitro* IFNα treatment, cells were stained with Sca-1-BV421. Stained cells were finally resuspended in staining media containing 1 µg/ml propidium iodide (PI) for dead cell exclusion. Cell sorting was performed on FACS Aria II SORP (Becton Dickinson) and each population was double sorted to maximize cell purity. Immunophenotyping was performed on either a Celesta (Becton Dickinson), ZE-5 (BioRad), or Novocyte Penteon (Agilent) cell analyzer. All data were analyzed with FlowJo (Treestar).

#### In vitro assays

All cultures were kept at 37°C in a 5% CO_2_ water jacket incubator (ThermoFisher Scientific) and all cytokines were purchased from PeproTech except IFNα (BioLegend) and IL-15 (ThermoFisher Scientific). For bulk liquid culture, sorted cells were grown in Iscove’s modified Dulbecco’s media (IMDM) containing 5% FBS (Gibco, 16140-071), 50 U/ml penicillin, 50 μg/ml streptomycin, 2mM L-glutamine, 0.1 mM non-essential amino acids, 1 mM sodium pyruvate and 50 μM 2-mercaptoethanol, EPO (4 U/ml), Flt3-L (25 ng/ml), GM-CSF (10 ng/ml), IL-3 (10 ng/ml), IL-11 (25 ng/ml), SCF (25 ng/ml), and TPO (25 ng/ml). The single cell differentiation culture assay was adapted from a previously published protocol^44^. Single cells were directly sorted into individual wells of a 96 well U-bottom plate in 200 µl of IMDM containing 10% FBS (Gibco, 16140-071), 20% BIT (StemCell Technology, 9500), 5% PFHM II (Gibco, 12040077), 50 U/ml penicillin, 50 µg/ml streptomycin, 2 mM L-glutamine, 0.1 mM non-essential amino acids, 1 mM sodium pyruvate, 55 µM 2-mercaptoethanol, EPO (4 U/ml), Flt3-L (10 ng/ml), GM-CSF (5ng/ml), IL-6 (10 ng/ml), SCF (50 ng/ml), and TPO (25 ng/ml). Wells containing megakaryocytes were manually scored; all other colonies were harvested and stained for flow cytometry after 6 to 8 days. The single cell OP9 differentiation culture assay was further adapted from another previously published protocol^45^. The day before the experiment, OP9 cells were plated at 1000-2000 cells/well onto a 0.1% gelatin-coated 96 well flat-bottom plate. The day of the experiment, the media was changed to 100 µl of differentiation media consisting of α−MEM containing 10% FBS, monothioglycerol (100 µM), ascorbic acid (50 µg/ml), 50 U/ml penicillin, 50 μg/ml streptomycin, and Glutamax (1X), supplemented with the following cytokines: EPO (4 U/ml), Flt3-L (10 ng/ml), IL-7 (10 ng/ml), IL-15 (5 ng/ml), SCF (10 ng/ml), and TPO (10 ng/ml). Single cells were directly sorted into individual wells. On day 5 of culture, cells were refreshed with 100 µl of differentiation media containing Flt3-L (10 ng/ml), IL-7 (10 ng/ml), and IL-15 (5 ng/ml). Wells containing megakaryocytes were manually scored; all other colonies were harvested and stained by flow cytometry after 10 days. For culture +/-IL-1β, cells were grown in StemPro34 medium (Gibco, 10639011) containing 50 U/ml penicillin, 50 μg/ml streptomycin, 2mM L-glutamine, EPO (4 U/ml), Flt3-L (25 ng/ml), GM-CSF (10 ng/ml), IL-3 (10 ng/ml), IL-11 (25 ng/ml), SCF (25 ng/ml), and TPO (25 ng/ml), with or without IL-1β (25 ng/ml). For stimulation with IFNα, cells were cultured in StemPro34 supplemented with SCF (25 ng/ml) and TPO (25 ng/ml) with or without IFNα (100 ng/ml) for 24 hours. For stimulation with IL-6 and pStat3 staining, c-Kit enriched cells were incubated at 37°C in 5% CO_2_ for 30 minutes in StemPro34 medium with or without IL-6 (50 ng/ml) and containing all the antibodies for HSPC staining (lineage cocktail, c-Kit-BV786, Sca-1-PE-Cy7, CD150-BV650, and CD48-AF700), then immediately washed, fixed, and permeabilized using the BD Phosflow Fix Buffer I and BD Phosflow Perm Buffer III according to the manufacturer’s instructions, and finally stained with pStat3-A647 for 1 hour at room temperature. Fixed cells were resuspended in staining media and analyzed on a ZE-5 (BioRad) cell analyzer.

#### Immunofluorescence staining

Cells (3,000-5,000 cells per slide) were pipetted onto poly-L-lysine coated slides (Sigma-Aldrich, P0425-72EA), incubated for 30 minutes at room temperature, fixed with 4% PFA for 10 minutes at room temperature, permeabilized with 0.3% TritonX-100/PBS for 2 minutes at room temperature and blocked in 1% BSA/PBS for 1 hour at room temperature. Slides were then incubated overnight at 4°C with rabbit anti-mouse p65, washed 3 times with PBS, incubated for 1 hour at room temperature in goat anti-rabbit IgG-A594, washed 3 times with PBS, incubated with 1 µg/ml DAPI for 5 minutes at room temperature, washed 2 times with PBS, and slides were mounted with ProLong Glass Antifade Mountant (ThermoScientific, P36980). Cells were imaged on a LeicaSP8 Upright Confocal Microscope (Leica, 40x objective) and images were processed using Imaris. At least 100 cells per condition were randomly captured for quantification. Nuclear vs. cytoplasmic p65 was scored by eye in a blinded manner.

#### Cytokine/chemokine analyses

Serum from maternal or fetal PB was collected by allowing whole blood to sit at room temperature for 20 minutes followed by centrifugation at 2,000 x g at 4°C. The supernatant was transferred to a new tube and frozen at −80°. Frozen serum was thawed and cytokine/chemokine levels were assessed using either the Mouse Inflammation Panel (BioLegend, 740446) or the Mouse Proinflammatory Chemokine Panel (BioLegend, 740451) according to the manufacturer’s instructions. Plates were run on a ZE-5 Cell Analyzer (BioRad). Data were analyzed using the LegendPlex Data Analysis Software Suite.

#### Quantitative RT-PCR

For qRT-PCR, 5,000-20,000 HSPCs were directly sorted into RLT Plus Lysis Buffer with 1% β-mercaptoethanol. RNA was extracted using a Direct-zol RNA Microprep Kit and reverse-transcribed using SuperScript IV VILO Master Mix. Runs were performed on QuantStudio 7 Flex Real-Time PCR System (Applied Biosystems) using SYBR Green and cDNA equivalents of 200 cells per reaction. Values were normalized to *Actb* expression.

#### Bulk RNA-sequencing

For bulk RNA-seq, RNA was isolated from 5,000-20,000 HSPCs using the RNeasy Plus Micro Kit. RNA integrity number (RIN) was determined by Bioanalyzer (Agilent Technologies) and RNA samples with RIN > 9.0 were further processed. RNA samples were processed over multiple experimental “sort days” and stored at −80°C in RNA isolation buffer until library preparation. A total of 3 biological replicates were sequenced for each population at each developmental stage. RNA was quantified and quality control (QC) was performed using an Agilent Bioanalyzer. Illumina sequencing libraries were prepared and sequenced at the NYU Genome Technology Center (New York, NY). The Automated NuGen Ovation Trio Low Input RNA-seq library preparation kit was used for library preparation, with all libraries prepared at the same time in order to minimize batch effects. Libraries were sequenced using an Illumina NovaSeq 6000 instrument with paired end 50 base pair sequencing to a target depth of 40 million unique reads per sample. Sequencing QC was performed using FastQC and MultiQC. Paired-end reads were mapped to the mouse reference genome (GRCm38 ver104) and counts were quantified using Salmon (v0.6.0)^83^. Gene counts were imported into a DDS object using DESeq2, removing genes with less than 100 reads summed across all included samples. The rlog transformation implemented in DESeq2 was utilized to normalize sample read counts for exploratory data analysis, including principal component analysis (PCA) and hierarchical clustering. The Benjamini-Hochberg algorithm was used for multiple testing correction, with a false discovery rate (FDR) = 0.05 significance threshold. Pathway analysis was performed for indicated pairwise comparisons using pre-ranked gene lists (meeting cut-off of log2 fold change (FC) > 1 and p adj < 0.05) on the Database for Annotation, Visualization, and Integrated Discovery (DAVID).

#### Single cell RNA sequencing

For scRNA-seq, 50,000 LK cells were sorted into 1.5 ml tubes containing 500 µl of HBSS with 2% FBS and transferred to the Columbia Genome Center Single Cell Analysis Core for microfluidic processing, library preparation and sequencing. Cells were re-counted, viability was assessed using a Countess II FL Automated cell counter (Thermo), and samples were processed following the manufacturer’s recommendations for Chromium Single Cell 3’ Library and Gel Bead Kit v.2 (10X Genomics). Samples were sequenced on an Illumina HiSeq4000, and alignment was performed using Cellranger (v.7.0.1) to mouse genome mm10. Downstream analyses were done using Seurat (v4)^86^ in R (V.3.1.1). QC was performed to exclude cells with the following parameters: < 200 genes, > 7000 genes, > 65,000 RNA counts, and > 7% mitochondrial genes; each gene had to be detected in at least 5 cells. Each sample was transformed using SCTransform before integration using the SCT method. G2M and S phase scores were assigned for each cell, and the difference between these scores regressed out to mitigate the heterogeneity of cell cycle stage while preserving the distinction between cycling and non-cycling cells^87^. After dataset integration, PCA selecting 30 principal components was performed and data visualized using UMAP. K parameter nearest neighbors were calculated before clusters were identified using the Louvain algorithm. Clusters were annotated using functions from the Semi-supervised Category Identification and Assignment (SCINA) R package^88^, using lists of genes for major cell types that were derived from integration of three independent ABM LK and LSK datasets using the FindConservedMarkers function in Seurat (Table S4). Each cluster was then named according to the prediction provided by SCINA of most likely identity, resulting in some variability among datasets as to precisely which cell types were represented by a discrete cluster. Clusters containing contaminating mature RBCs and lymphocytes were removed from the analysis. Differentially expressed genes (DEG) were identified within defined clusters using the FindMarkers function (min % 0.25 and log2FC threshold 0.25). gene set enrichment analysis (GSEA) was performed using the ClusterProfiler R package^89,90^ using the gseGO function with the following parameters: minimum gene set size 10, maximum gene set size 800, p value < 0.05. Gene lists for module scores were downloaded from gene sets in MSigDB and added using the AddModuleScore function. Violin plots were generated using the VlnPlot function.

#### Single cell ATAC sequencing

For scATAC-seq, 50,000 LK or LSK cells were sorted into 1.5 ml tubes containing 500 µl of HBSS with 2% FBS and allowed to rest on ice for 1 hour prior to centrifugation. Cells were pelleted at 350 x g for 5 minutes at 4°C and resuspended in 0.04% BSA in PBS and pelleted again at 350 x g for 5 minutes at 4°C. Supernatant was removed, and the pelleted cells were lysed by adding 45 µL of chilled lysis buffer (10mM Tris-HCl (pH 7.4), 10mM NaCl, 3mM MgCl2, 0.1% Tween-20, 0.1% NP-40 Substitute, 0.01% Digitonin, 1% BSA, 1mM DTT, 1U/µL RNase inhibitor) and pipetted gently 3 times before being incubated on ice for 3 minutes. Resulting nuclei were washed by adding 50 µL of chilled wash buffer (10mM Tris-HCl (pH 7.4), 10mM NaCl, 3mM MgCl2, 0.1% Tween-20, 1% BSA, 1mM DTT, 1U/µL RNase inhibitor). Nuclei were pelleted at 500 x g for 10 minutes at 4°C and were washed in diluted nuclei buffer (10X Genomics). Multiome GEM generation and library preparation was performed according to 10X Genomics protocol CG000338, targeting 5000 cell data recovery. ATAC libraries were pooled 1:1:1:etc, sequenced on an Illumina NovaSeq 5000, and alignment was performed using Cellranger (v.7.0.1) to mouse genome mm10.. Gene Expression libraries were pooled 1:1:1:etc, sequenced on an Illumina NovaSeq 5000, and alignment was performed using Cellranger (v.7.0.1) to mouse genome mm10. The single cell RNA-seq data did not pass the QC parameter of % mitochondrial reads and was therefore not used for further analysis. The single cell ATAC-seq data was analyzed using Signac^91^ with the following QC parameters used to filter poor quality cells: ATAC count > 1000 and < 100000 per cell, nucleosome signal < 2, TSS enrichment > 1. Samples were integrated using the IntegrateEmbeddings function. A combined peaks file was generated for all 4 component datasets by merging peak files that were generated using Macs2^92^ for each component dataset; this combined peak file was used to annotate peaks in each component dataset prior to integration. After dataset integration, PCA selecting 30 principal components was performed and data visualized using UMAP, using component 2 through 30. K parameter nearest neighbors were calculated and clusters identified using the smart local moving (SLM) algorithm. Clusters were manually annotated based on DEGs from the RNA assay. Differentially accessible peaks were evaluated using the FindMarkers function with the logistic regression test with a latent variable of the total peak count per cell, with peaks that were detectable in minimum 5% of cells and log2FC threshold 0.1. GSEA was performed using the ClusterProfiler R package using the gseGO function using the gene closest to differentially accessible peaks, with the following parameters: minimum gene set size 10, maximum gene set size 800, p value < 0.05. Transcription factor (TF) motif enrichment analysis was performed using the FindMotifs function and the JASPAR motif database^93^. TF footprinting was performed using the Footprint function for indicated TF motifs.

### Quantification and Statistical Analysis

Data are represented as means ± standard deviations (S.D.), or as violin plots with the center line representing the median, using R studio for RNA-seq and ATAC-seq data or GraphPad Prism (V.9.5.0) for all other data. Circles on bar graphs represent biological replicates. Student t-test was used when two groups were compared, with Holm-Šídák correction when multiple comparisons were performed, one-way ANOVA when three or more groups were compared, with Bonferroni’s, Dunnett’s, or Tukey’s correction when multiple comparisons were performed, and Mantel-Cox log rank test for comparison of fetus^WT-mom^ vs. fetus^KO-mom^ mice survival.

