## Supplemental Figures for "Maternal IL-10 restricts fetal emergency myelopoiesis"

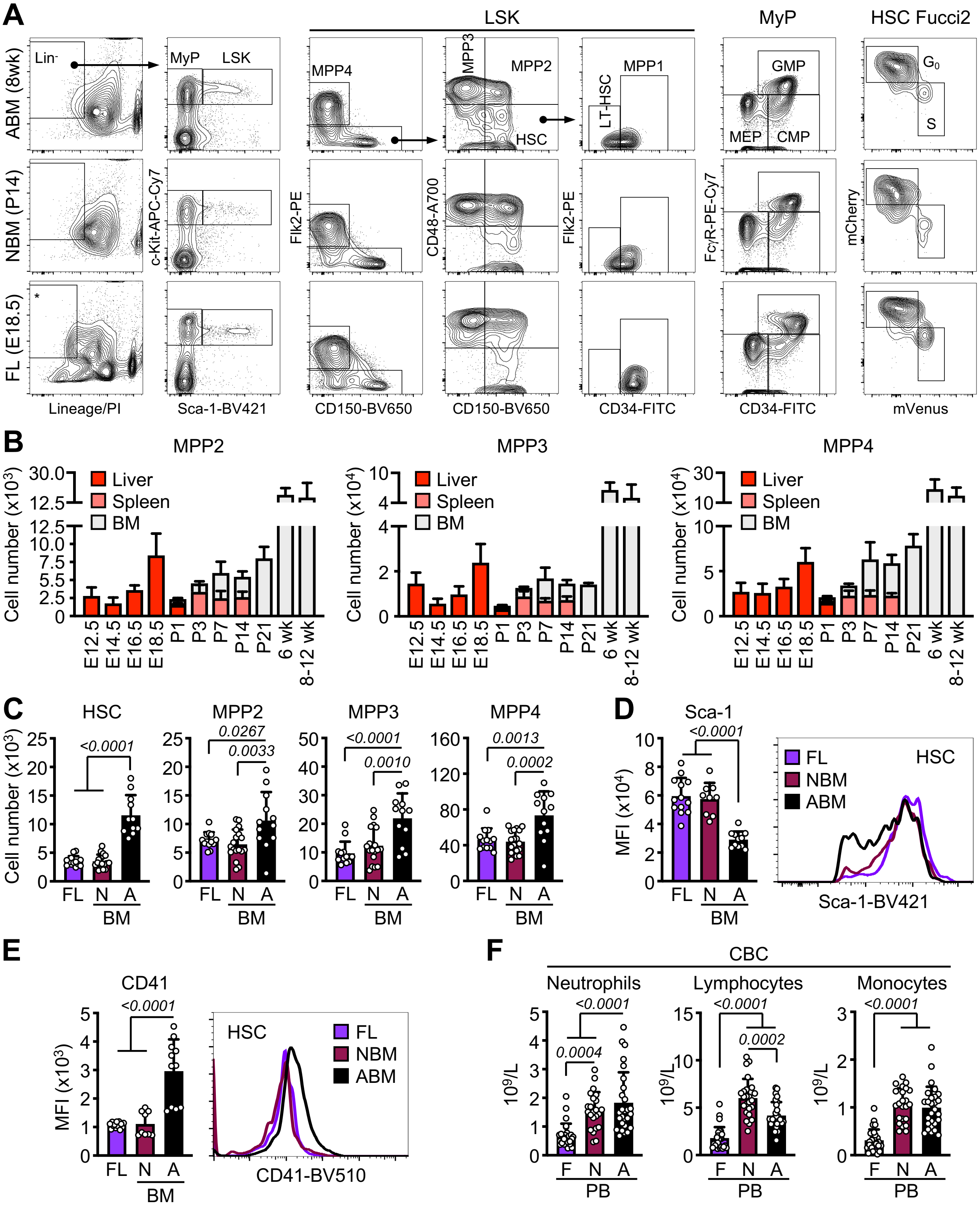


**Figure S1** (Collins et al.)

**Figure S1. Decreased ground state myelopoiesis during fetal to adult transition, related to Figure 1.**
**A**) Representative flow cytometry plots and gating strategy for HSPC subsets in LSK cells, myeloid progenitors in MyP cells, and cell cycle distribution in HSCs from Fucci2 cell cycle reporter mice in fetal liver (FL), neonatal BM (NBM), and adult BM (ABM). * Mac-1 and CD4 were omitted from the lineage (Lin) cocktail for FL analyses.
**B**) MPP2 (left), MPP3 (middle), and MPP4 (right) number in liver, spleen, or bone marrow (BM) of fetuses (F), E12.5 (6), E14.5 (13), E16.5 (7), E18.5 (8), neonates (N), P1 (4), P3 (4), P7 (9), P14 (5), and P21 (5), and adult (A), 6wk (6) and 8-12wk (7), mice; wk, week. Results are from the number of biological repeat (individual mice) from at least 2 independent experiments.
**C**) Number of each HSPC population in E18.5 FL, P14 NBM, and 8wk ABM (3 independent experiments).
**D**) Median fluorescence intensity (MFI) (left) and representative histogram (right) of Sca-1 staining in FL, NBM, and ABM HSCs (3 independent experiments).
**E**) MFI (left) and representative histogram (right) of CD41 staining in FL, NBM, and ABM HSCs (3 independent experiments).
**F**) Number of neutrophils (left), lymphocytes (middle), and monocytes (right) from complete blood counts (CBC) of peripheral blood (PB) from fetal, neonatal, and adult mice (5 independent experiments).
Data are means ± S.D. Circles represent biological replicates (individual mice).

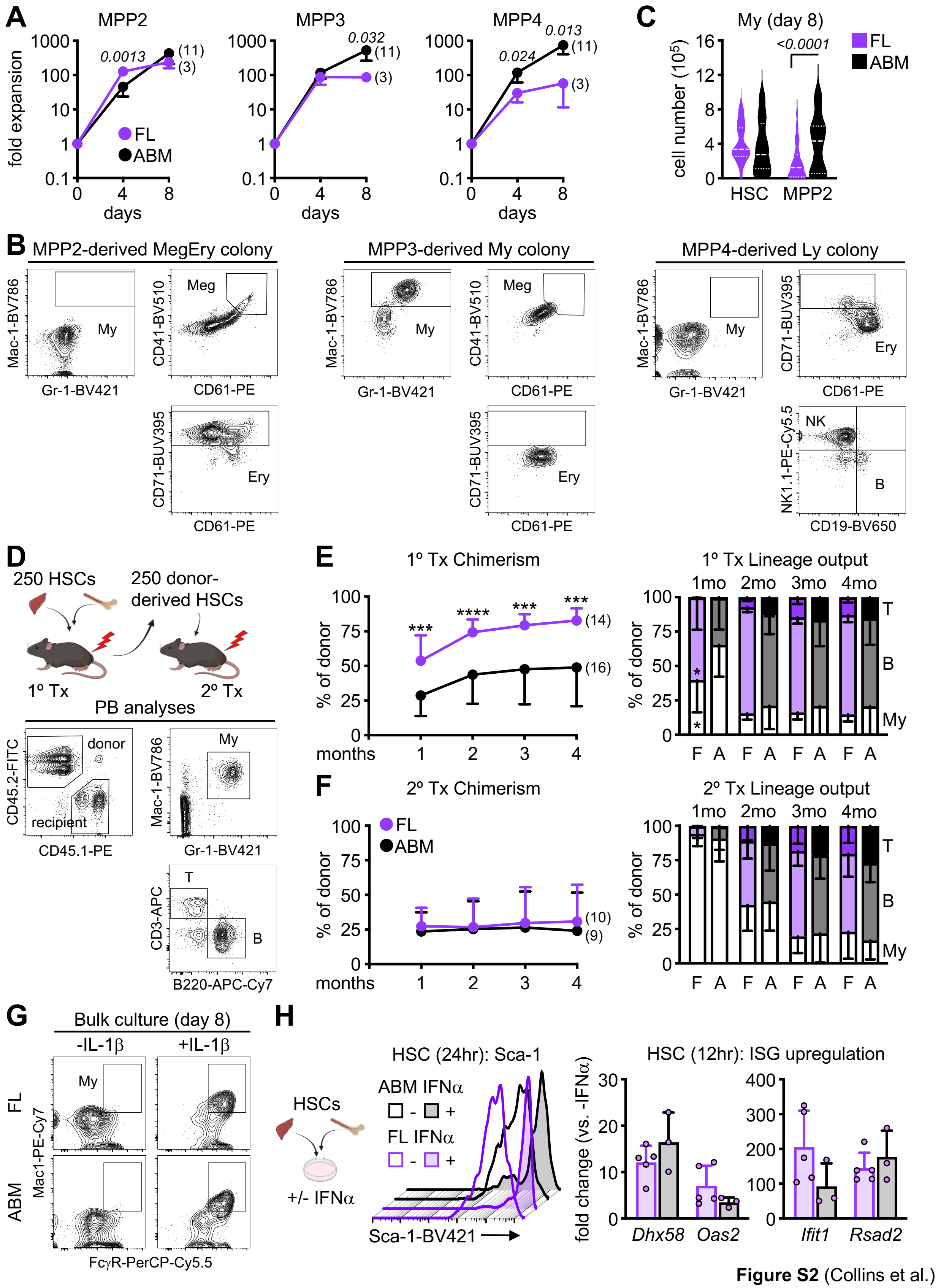


**Figure S2** (Collins et al.)

**Figure S2. Extended characterization of fetal HSPC, related to Figure 2.**

**A**) Fold expansion of FL and ABM MPPs in bulk liquid culture (3 independent experiments).

**B**) Representative flow cytometry plots and gating strategy for *in vitro* single cell HSPC differentiation culture with examples of day 6 MPP2-derived megakaryocyte/erythroid (MegEry) colony (left) and MPP3-derived myeloid (My) colony (middle), and day 10 MPP4-derived lymphoid (Ly) colony (right).

**C**) Cellularity of myeloid (My) colonies at day 8 of single cell differentiation culture of FL and ABM HSC and MPP2 (2 independent experiments with a range of 21-73 colonies counted per population). Results are shown as violin plots with median and quartiles.

**D**) Schematic for primary (1°) and secondary (2°) transplantations of FL and ABM HSCs in lethally irradiated recipients with representative flow cytometry plots of gating strategy for chimerism and lineage output; My, myeloid; B, B cells; T, T cells.

**E-F**) Donor chimerism (left) and donor-derived lineage output (right) over time in the peripheral blood of primary (E) and secondary (F) recipients (2 independent experiments); F, FL; A, ABM; mo, month.

**G**) Representative flow cytometry plots of myeloid (My) cell staining after 8 days of FL and ABM HSCs culture with or without (+/-) IL1b (25 ng/ml).

**H**) Schematic of bulk liquid culture of FL and ABM HSCs +/- IFNa (100 ng/ml) (left) with representative FACS histograms of Sca-1 staining upon 24 hours (hr) culture (middle), and qRT-PCR of representative interferon-stimulated genes (ISG) upon 12 hours culture (right) (3 independent experiments).

Data are means ± S.D. except for (C); $*p_{adj} \leq 0.05$; $***p_{adj} \leq 0.001$; $****p_{adj} \leq 0.0001$. Number of biological replicates and transplanted mice are indicated either in parentheses or with individual circles.

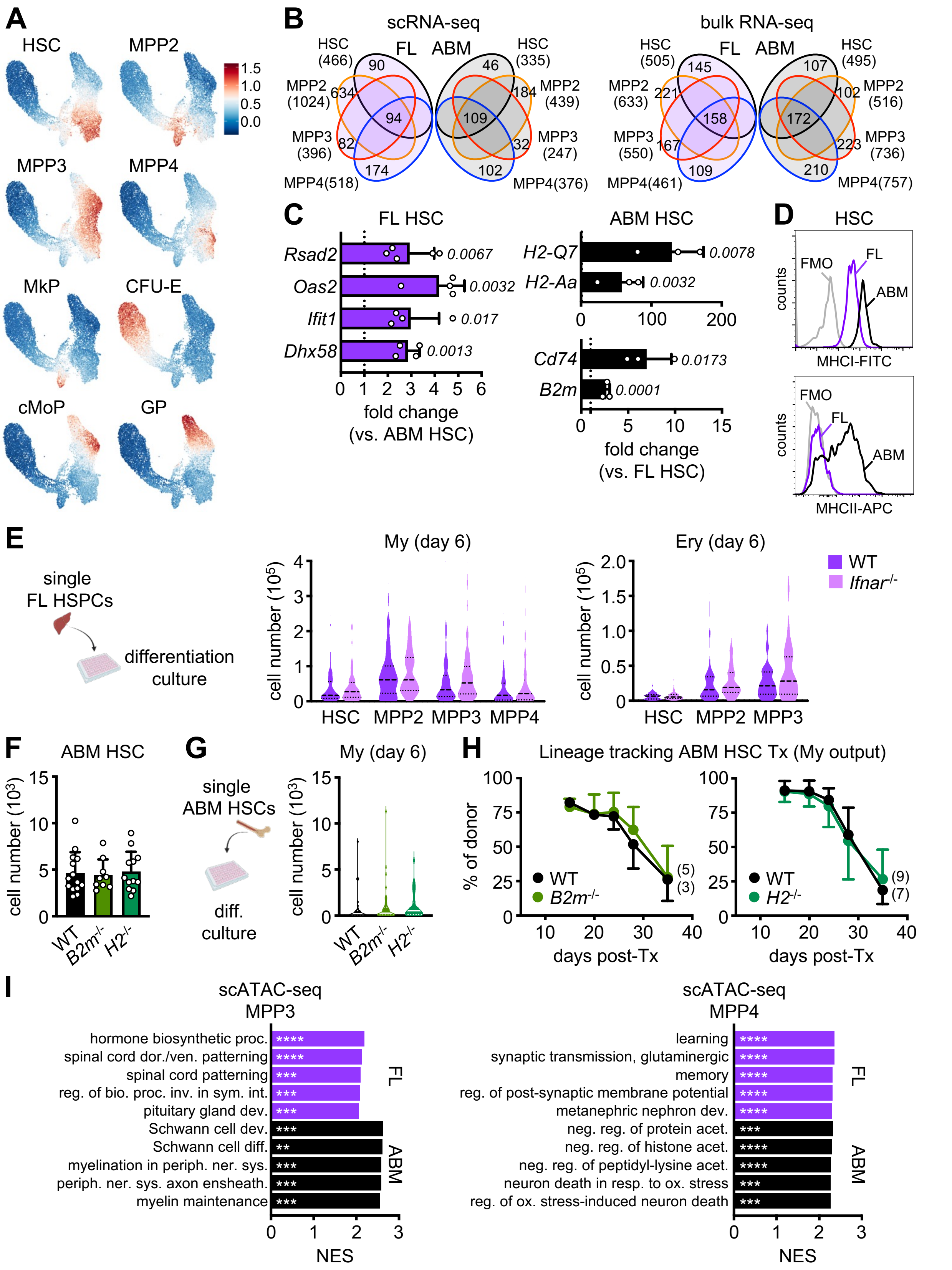


**Figure S3** (Collins et al.)

**Figure S3. Detailed characterization of fetal and adult HSPC transcriptional signatures, related to Figure 3.**

**A**) Projection of module score for cluster identification using manually curated ID genes.

**B**) Venn diagrams showing the number of unique and shared upregulated differentially enriched genes (DEG) found in FL and ABM HSPCs in scRNA-seq (left) and bulk RNA-seq (right) datasets.

**C**) qRT-PCR validation of signature interferon-stimulated genes (ISG) in FL HSCs (left) and antigen processing and presentation genes in ABM HSCs (3-5 independent experiments).

**D**) Representative FACS histograms of MHCI and MHCII expression on FL and ABM HSCs; FMO, fluorescence minus one.

**E**) Schematic of single cell differentiation culture of FL wild type (WT) and *Ifnar*$^{-/-}$ HSPCs with cellularity of myeloid (My, left) and erythroid (Ery, right) colonies at day 8 (3 independent experiments with a range of 26-132 colonies counted per population). Cellularity data are shown as violin plots with median and quartiles.

**F**) Quantification of HSCs in adult WT, MHCI-deficient *B2m*$^{-/-}$, and MHCII-deficient *H2*$^{-/-}$ mice (3 independent experiments).

**G**) Schematic of single cell differentiation culture of ABM WT, *B2m*$^{-/-}$, and *H2*$^{-/-}$ HSCs with cellularity of myeloid (My) colonies at day 8 (3 independent experiments with a range of 28-44 colonies counted per population). Cellularity data are shown as violin plots with median and quartiles.

**H**) Lineage tracking transplantation (Tx) of ABM WT, *B2m*$^{-/-}$, and *H2*$^{-/-}$ HSCs in sub-lethally irradiated recipients with percent donor-derived myeloid output in peripheral blood at the indicated days post-transplantation (2 independent experiments).

**K**) Top 5 pathways driven by genes closest to open chromatin regions (OCR) found differentially enriched in MPP3 (left) and MPP4 (right) clusters from FL and ABM scATAC-seq (GSEA on differentially accessible ORCs); proc., process; dor./ven., dorsal/ventral; reg., regulation; bio., biological; inv., involved; sym., symbiotic; int., interaction; dev., development; diff., differentiation; periph., peripheral; ner., nervous; sys., system; ensheath., ensheathment; neg., negative; acet., acetylation; resp., response; ox., oxidative.

Data are means ± S.D except for (E and G). ** $p_{adj} \leq 0.01$; ***$p_{adj} \leq 0.001$; ****$p_{adj} \leq 0.0001$. Number of biological replicates and transplanted mice are indicated either in parentheses or with individual circles.

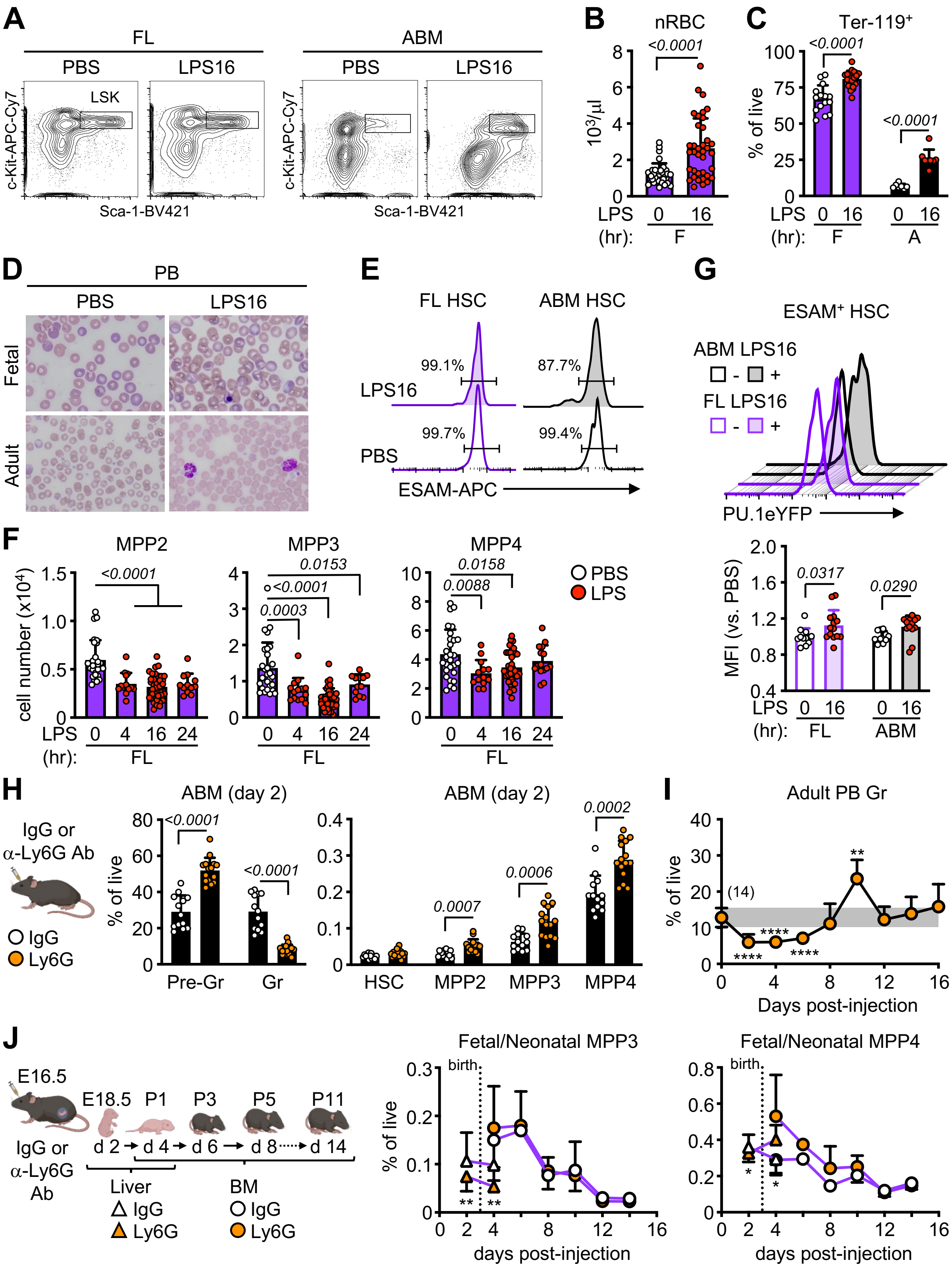


**Figure S4** (Collins et al.)

**Figure S4. Lack of emergency myelopoiesis engagement in fetal HSPCs, related to Figure 4.**
**A**) Representative flow cytometry plots of LSK cells in LPS-exposed FL and ABM cells showing Sca-1 upregulation only in adult mice.
**B**) Nucleated RBC (nRBC) count from CBC of LPS-exposed fetal PB (5 independent experiments).
**C**) Quantification of Ter119$^{+}$ cells in PB from LPS-exposed fetal and adult mice (3 independent experiments).
**D**) Representative Wright-Giemsa staining of PB smears of LPS-exposed fetal and adult mice.
**E**) Representative flow cytometry histograms of ESAM staining on LPS-exposed FL and ABM HSCs.
**F**) Quantification of MPP2 (left), MPP3 (middle), and MPP4 (right) in LPS-exposed FL (5 independent experiments).
**G**) Representative FACS histograms (top) and quantification (bottom) of PU.1-eYFP MFI in LPS-exposed FL and ABM ESAM$^{+}$ HSCs (3 independent experiments in *PU.1-eYFP* reporter mice).
**H**) Schematic of IgG or anti-Ly6G antibody (Ab) adult injection (1 mg/mouse) with quantification of myeloid cells (left) and HSPCs (right) in ABM 2 days after exposure (4 independent experiments); Pre-Gr, pre-granulocyte (Mac-1$^{+}$/Gr-1$^{int}$); Gr, granulocyte (Mac-1$^{+}$/Gr-1$^{+}$).
**I**) Quantification of granulocytes in PB of Ly6G-exposed adult mice (2 independent experiments). Grey shade indicates the range steady state granulocyte level.
**J**) Schematic of IgG or anti-Ly6G antibody fetal injection (1 mg/mouse) with quantification of MPP3 (left) and MPP4 (right) in liver (triangles) and BM (circles) of fetal/neonatal mice exposed to IgG or Ly6G during development (3 independent experiments; see Table S1). Dotted line indicates birth.
Data are means ± S.D.; *$p_{adj} \leq 0.05$; ** $p_{adj} \leq 0.01$; ****$p_{adj} \leq 0.0001$. Number of biological replicates and treated fetuses/mice are indicated either in parentheses or with individual circles.

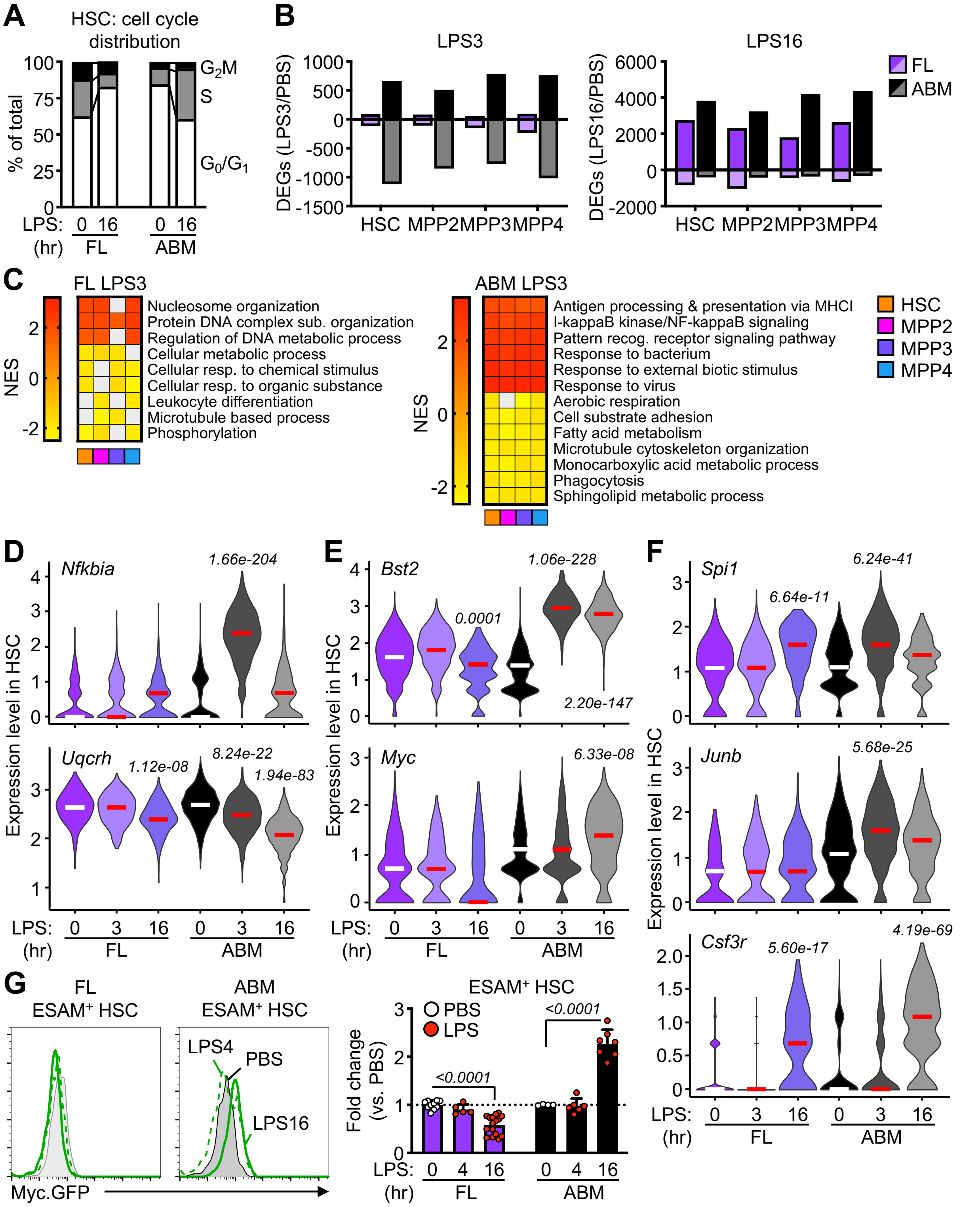


**Figure S5** (Collins et al.)

**Figure S5. Lack of emergency myelopoiesis engagement in fetal HSPCs at the molecular level, related to Figure 5.**

**A**) Cell cycle distribution in HSC cluster from FL and ABM LK cells exposed to PBS or LPS for 16 hours.

**B**) Total number of DEGs in HSPC clusters from FL and ABM LK cells isolated from mice exposed to LPS for 3 (left) or 16 (right) hours.

**C**) Top pathways differentially enriched in FL (left) and ABM (right) HSPC clusters from 3 hours LPS-exposed FL and ABM scRNA-seq (GSEA on DEGs; $\log_2$ FC ± 0.25, min.pct 0.25). Grey boxes are pathways not meeting the $p_{adj} < 0.05$ cut-off.

**D**) *Nfkbia* and *Uqcrh* expression in HSC cluster from LPS-exposed FL and ABM scRNA-seq.

**E**) *Bst2* and *Myc* expression in HSC cluster from 3 hours LPS-exposed FL and ABM scRNA-seq.

**F**) *Spi1, Junb*, and *Csf3r* expression in HSC cluster from 3 hours LPS-exposed FL and ABM scRNA-seq.

**G**) Representative FACS histograms (left) and quantification (right) of GFP MFI on LPS-exposed ESAM$^{+}$ HSCs from Myc.GFP reporter mice (5 independent experiments).

Data are means ± S.D. Number of biological replicates and treated fetuses/mice are indicated with individual circles.

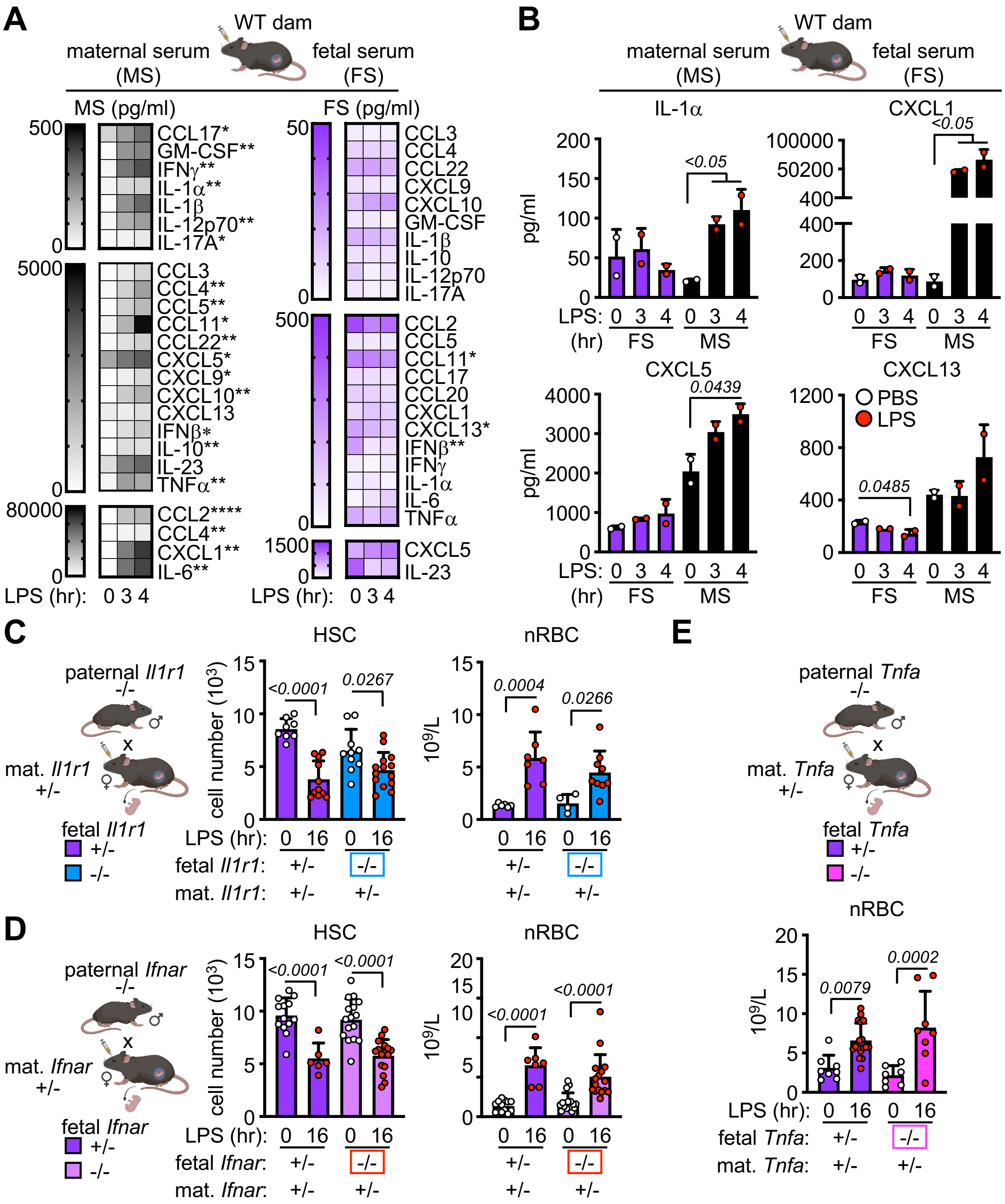


Figure S6 (Collins et al.)

**Figure S6. Extrinsic regulation of fetal emergency myelopoiesis, related to Figure 6.**
**A**) Cytokine levels in paired maternal serum (left, MS) and fetal serum (right, FS) from mice exposed to LPS for 3 and 4 hours. Color intensity represents the mean of 2 technical replicates (2 independent experiments). Fetal serum is pooled from all fetuses in the litter (average 5 to 8 fetuses per litter).
**B**) Levels of the indicated cytokines in paired fetal and maternal serum from LPS-exposed mice.
**C**) Schematic of *Il1r1* time-mated pregnancies with quantification of FL HSCs (left) and PB nucleated RBCs (nRBC, right) in LPS-exposed *Il1r1*$^{+/-}$ and *Il1r1*$^{-/-}$ fetuses (3 independent experiments).
**D**) Schematic of *Ifnar* time-mated pregnancies with quantification of FL HSCs (left) and PB nRBCs (right) in LPS-exposed *Ifnar*$^{+/-}$ and *Ifnar*$^{-/-}$ fetuses (4 independent experiments).
**E**) Schematic of *Tnfa* time-mated pregnancies with quantification of PB nRBCs in LPS-exposed *Tnfa*$^{+/-}$ and *Tnfa*$^{-/-}$ FL (4 independent experiments).
Data are means ± S.D.; *$p_{adj} \leq 0.05$; **$p_{adj} \leq 0.01$; ****$p_{adj} \leq 0.0001$. Number of biological replicates and treated fetuses/mice are indicated with individual circles.

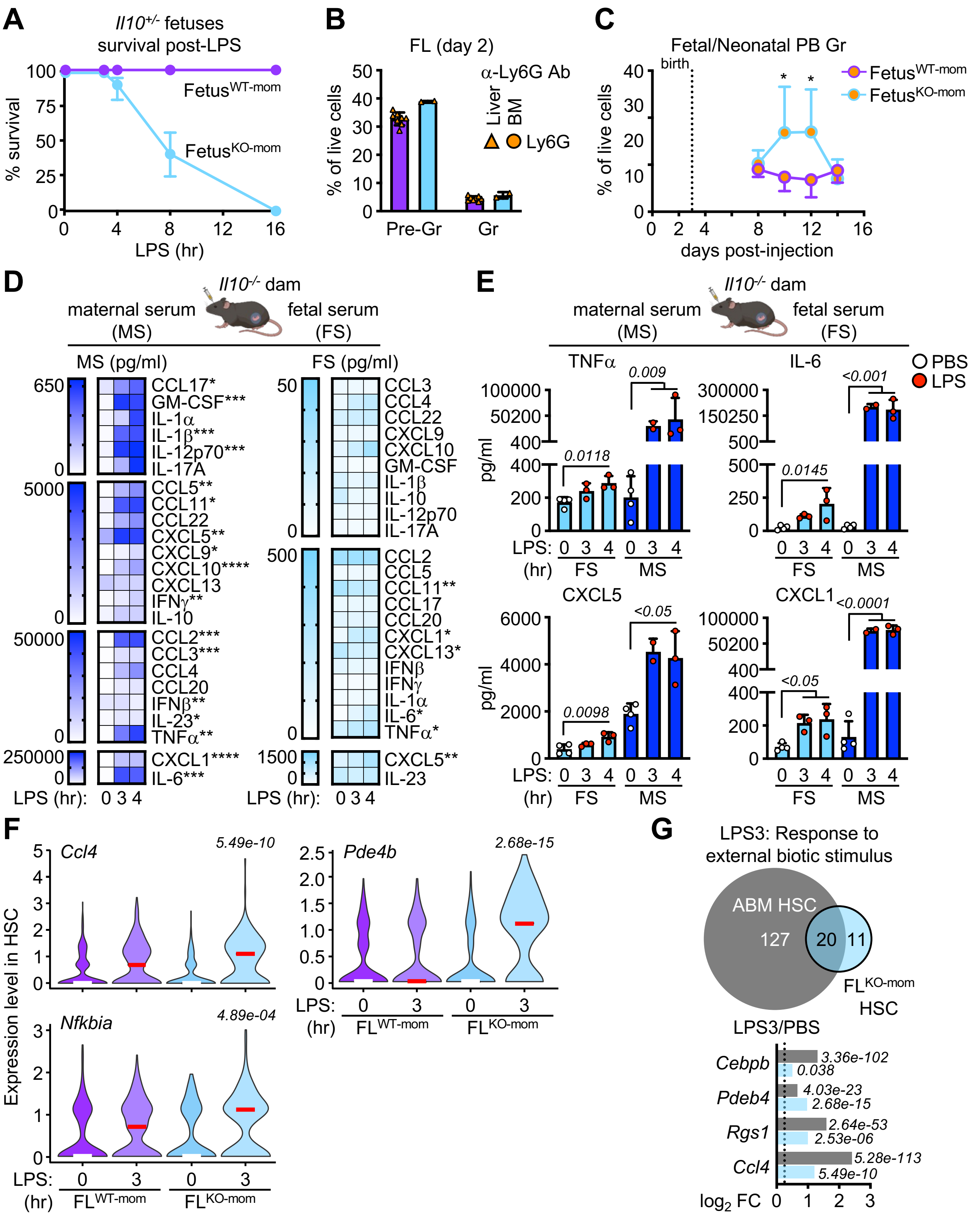


**Figure S7** (Collins et al.)

**Figure S7. Maternal IL-10 restricts fetal emergency myelopoiesis, related to Figure 7.**
**A**) Survival of *Il10*$^{-/-}$ fetuses in fetus$^{WT\text{-}mom}$ and fetus$^{KO\text{-}mom}$ mice in response to LPS (at least 2 independent experiments; see Table S2).
**B**) Quantification of FL myeloid cells 2 days after Ly6G exposure in fetus$^{WT\text{-}mom}$ and fetus$^{KO\text{-}mom}$ mice (2 independent experiments); Pre-Gr, pre-granulocyte (Mac-1$^{+}$/Gr-1$^{int}$); Gr, granulocyte (Mac-1$^{+}$/Gr-1$^{+}$).
**C**) Quantification of granulocytes in the PB of fetal/neonatal fetus$^{WT\text{-}mom}$ and fetus$^{KO\text{-}mom}$ mice exposed to a-Ly6G during development (at least 2 independent experiments; see Table S3). Dotted line indicates birth.
**D**) Cytokine levels in paired maternal serum (left, MS) and fetal serum (right, FS) from fetus$^{KO\text{-}mom}$ mice exposed to LPS for 3 and 4 hours. Color intensity represents the mean of 2 technical replicates (4 independent experiments). Fetal serum is pooled from all fetuses in the litter (average 5 to 8 fetuses per litter).
**E**) Levels of the indicated cytokines in paired fetal and maternal serum from LPS-exposed fetus$^{KO\text{-}mom}$ mice.
**F**) *Ccl4*, *Pde4b*, and *Nfkbia* expression in HSC cluster from 3 hours LPS-exposed fetus$^{WT\text{-}mom}$ and fetus$^{KO\text{-}mom}$ FL scRNA-seq.
**G**) Number of unique and shared upregulated DEGs in HSC cluster from ABM scRNA-seq and HSC cluster from fetus$^{KO\text{-}mom}$ FL scRNA-seq. Bar graph shows $\log_2$FC for selected genes.
Data are means ± S.D.; *$p_{adj} \leq 0.05$; **$p_{adj} \leq 0.01$; ***$p_{adj} \leq 0.001$; ****$p_{adj} \leq 0.0001$. Number of biological replicates and treated fetuses/mice are indicated with individual circles.

**Table S1. Biological replicates for fetal/neonatal Ly6G exposure kinetics in WT mice, related to Figures 4J and S4J.**

| Sample | IgG | anti-Ly6G |
|---|---|---|
| Day 2 FL | 25 | 36 |
| Day 4 FL | 6 | 8 |
| Day 4 BM | 2* (4; 3) | 3* (4; 3; 3) |
| Day 6 BM | 2* (3; 2) | 2* (6; 3) |
| Day 8 BM | 3* (3; 2; 3) | 4* (4; 3; 3; 3) |
| Day 10 BM | 7 | 8 |
| Day 12 BM | 8 | 7 |
| Day 14 BM | 6 | 7 |

Data represent the number of individual fetuses or neonatal mice collected at each time point following IgG or anti-Ly6G treatment of WT dams except when indicated. * indicates when individual biological replicates from pooled neonatal mice was used instead, with the number of pups per replicate provided in parentheses.

**Table S2. Biological replicates for fetus survival post-LPS exposure in fetus$^{WT\text{-}mom}$ and fetus$^{KO\text{-}mom}$ mice, related to Figure S7A.**

| Hours of LPS | Fetus$^{WT\text{-}mom}$ | Fetus$^{KO\text{-}mom}$ |
|---|---|---|
| 0 | (22/22); 3 | (32/32); 5 |
| 3 | | (13/13); 2 |
| 4 | (25/25); 4 | (22/27); 5 |
| 8 | | (0/17); 2 |
| 16 | (17/17); 2 | (0/14); 2 |

Data in parentheses represent the number of surviving fetuses/total fetuses recovered from the indicated number of pregnant dams.

**Table S3. Biological replicates for fetal/neonatal Ly6G exposure kinetics in fetus$^{WT\text{-}mom}$ and fetus$^{KO\text{-}mom}$ mice, related to Figures 7E and S7C.**

| Sample | Fetus$^{WT\text{-}mom}$ | Fetus$^{KO\text{-}mom}$ |
|---|---|---|
| Day 2 FL | 8 | 2 |
| Day 8 BM/PB | 6 | 16 |
| Day 10 BM/PB | 6 | 9 |
| Day 12 BM/PB | 6 | 12 |
| Day 14 BM/PB | 4 | 7 |

Data represent the number of individual fetuses or neonatal mice collected at each time point following anti-Ly6G treatment of *Il10*-competent (fetus$^{WT\text{-}mom}$) and *Il10*-deficient (fetus$^{KO\text{-}mom}$) dams.

**Table S4. Gene lists for cluster identification, related to star methods.**

| LT-HSC | ST-HSC | MPP2 | MPP3 | MPP4 | MkP |
|---|---|---|---|---|---|
| *Hlf* | *Ctla2a* | *Apoe* | *H2afy* | *Wfdc17* | *Cavin2* |
| *Ltb* | *Ifitm1* | *Itga2b* | *Emb* | *Ighm* | *Pbx1* |
| *Ctla2a* | *Pim1* | *Pbx1* | *Ccl9* | *Dntt* | *Pf4* |
| *Ifitm1* | *Adgrl4* | *Gata2* | *Cd34* | *Flt3* | *Gp9* |
| *Mecom* | *Hlf* | *F2r* | *Irf8* | *H2afy* | *Tuba8* |
| *Procr* | *Adgrg1* | *Ctla2a* | *BC035044* | *Lsp1* | *Cd9* |
| *Txnip* | *Gcnt2* | *Nrgn* | *Ms4a6c* | *Il12a* | *Vwf* |
| *Gcnt2* | *Sox4* | *Fgf3* | *Bex6* | *AA467197* | *Apoe* |
| *Gimap1* | *Gm19590* | *Zfpm1* | *Psmb8* | *Satb1* | *Nrgn* |
| *Rgs1* | *Tespa1* | *Pdzk1ip1* | *Plac8* | *Egfl7* | *Itga2b* |
| *Pdzk1ip1* | *Cd34* | *Sdsl* | *Bcl2* | *Cox6a2* | *F2r* |
| *Gm19590* | *Zfp36l2* | *Rpl21* | *Tmsb10* | *Cmah* | *F2rl2* |
| *Pnrc1* | *Ltb* | *Angpt1* | *Igfbp4* | *Mef2c* | *Gp1bb* |
| *Malat1* | *Txnip* | *Fermt3* | *Glipr1* | *Ccl3* | *Gp5* |
| *Gimap6* | *Gimap1* | *Rgs1* | *Cd48* | *Gm5111* | *Treml1* |
| *Rbp1* | *Ptpn18* | *Meis1* | *Sell* | *Myl10* | *Rab27b* |
| *Angpt1* | *Tmem176b* | *Gp9* | *Ramp1* | *Notch1* | *Gata2* |
| *Mpl* | *Lmo2* | *Unc119* | *H2-DMa* | *Cd34* | *Fermt3* |
| *Msi2* | *Ifi203* | *Cd9* | *Lat2* | *Arpp21* | *Unc119* |
| *Nkx2-3* | *Ptpre* | *Dapp1* | *Satb1* | *Tcf4* | *Zfpm1* |
| *Ifitm3* | *Ifitm3* | *Hmgb3* | *Pou2f2* | *Gpr171* | *Mfsd2b* |
| *Meis1* | *Ptprcap* | *Vamp5* | *Fkbp1a* | *Gimap6* | *Ctla2a* |
| *Myct1* | *Vldlr* | *Gp5* | *Bmyc* | *Cd33* | *Angpt1* |
| *Ly6a* | *Msi2* | *Smim5* | *Tespa1* | *Ptprcap* | *Mpl* |
| *Shisa5* | *Ly6a* | *Mfsd2b* | *Ly86* | *Cd52* | *Meis1* |
| *Hacd4* | *Akap13* | *Slc18a2* | *Cd93* | *H2-Ob* | *Rpl21* |
| *Lmo2* | *Gimap6* | *Diaph1* | *Fam117a* | *Ifi27l2a* | *Slc14a1* |
| *Zfp36l2* | *Angpt1* | *Slc22a3* | *Fabp5* | *Sox4* | *Cuedc1* |
| *Adgrg1* | *Pnrc1* | *Mpl* | *Adgrg3* | *Tbxa2r* | *Plek* |
| *Gimap5* | *Myl10* | *Rab37* | *Hist1h2bc* | *Lck* | *Dapp1* |
| *Myl10* | *Gm5111* | *Adgrg1* | | *Tespa1* | *Rgs18* |
| *Zbtb20* | *Cd27* | *Gng11* | | *Sdc4* | *Gng11* |
| *Pim1* | *Rbpms* | *Gnb4* | | *Ifi203* | *Fyb* |
| *Pik3ip1* | *Dapp1* | *Fyb* | | *Wfdc18* | *Mef2c* |
| *Adgrl4* | *Meis1* | *Plxnc1* | | *Tsc22d1* | *Vamp5* |
| *Tsc22d3* | *Shisa5* | *Serpina3g* | | *Ppp1r18* | *Smim5* |
| *Rbm39* | *Malat1* | *Tspan32* | | *Laptm5* | *Rab37* |
| *Gng11* | *Paip2* | *Trim47* | | *Sstr2* | *Lat* |

| | | | | | |
|---|---|---|---|---|---|
| Akap13 | Zyx | Rgs18 | | Cd27 | Hmgb3 |
| Ifi203 | Myct1 | Mef2c | | Pou2f2 | Rap1b |
| Tmem176b | Ypel3 | H2-Q7 | | Tmsb10 | Pdzk1ip1 |
| Glul | B2m | Cbfa2t3 | | Ramp1 | Fgf3 |
| Ptprcap | Ikzf2 | Muc13 | | Shisa8 | Cavin3 |
| H2-K1 | AC149090.1 | Tgfb1 | | Dusp2 | Serpina3g |
| B2m | Tsc22d1 | Nt5c3 | | Tmem108 | Tgfb1 |
| Esam | Lsp1 | Bin2 | | Smim14 | Diaph1 |
| AC149090.1 | Laptm5 | Lat | | Il7r | Plxnc1 |
| Cdkn1c | Wfdc17 | Gata1 | | Lat2 | Rabgap1l |
| Kit | Rbm39 | Gse1 | | Marcksl1 | Nt5c3 |
| Trim47 | Cox7a2l | Ddah2 | | Pim1 | Tmem40 |
| **BaP** | **Pre-CFU-E** | **CFU-E** | **GMP_mult** | **cMoP** | **GP** |
| Ms4a2 | Ptpn14 | Bag2 | Fcgr3 | Fcer1g | Fcgr3 |
| Csrp3 | Optn | Hs6st1 | Hspa5 | Ifi209 | Hsd11b1 |
| Cpa3 | Car1 | Cox5b | Hdc | Slpi | G0s2 |
| Cyp11a1 | Gstm5 | Mrpl30 | Cst7 | Ctsz | Fcnb |
| Gm11697 | Ugcg | Il1rl1 | Slpi | Tctex1d1 | S100a8 |
| Alox5 | Ermap | Gm37915 | Gstm1 | AI839979 | Sgms2 |
| Fcer1a | Atpif1 | Slc40a1 | Sec61b | Anxa3 | Chd7 |
| Csf1 | Abcb4 | Hspd1 | Anxa3 | Plac8 | Prom1 |
| Gzmb | Aqp1 | Hspe1 | Plac8 | Erp29 | Cldn15 |
| Fam129a | Vamp5 | Orc2 | Cldn15 | Met | Mgam |
| Rnase12 | Apoe | Abcb6 | Clec12a | Clec5a | Chst13 |
| Fgf3 | Csrp3 | Ncl | Pglyrp1 | Rassf4 | Clec4a2 |
| 4930519L02Rik | Klf1 | Steap3 | Dmkn | Clec12a | Far2 |
| St8sia6 | Ces2g | Ctse | Nkg7 | Mgst1 | Pglyrp1 |
| Slc45a3 | Gm15915 | Camsap2 | Igsf6 | Tyrobp | Dmkn |
| Gata2 | Mfsd2b | Mgst3 | Calr | Igsf6 | Mogat2 |
| Lmo4 | Trib2 | Ppox | Hp | Nrg1 | Nucb2 |
| Slc18a2 | Tspo2 | Spta1 | Alas1 | Cyp4f18 | Igsf6 |
| Scin | Kcng2 | Ahctf1 | Manf | Hp | Mcemp1 |
| Ikzf2 | Blvrb | Psen2 | Prss57 | Irf8 | Ncam1 |
| Cd63 | Atp1b2 | Lefty1 | Prtn3 | Nrp1 | Mgl2 |
| Hdc | Zg16 | Optn | Elane | Ctsh | Rflnb |
| Lpcat2 | Samd14 | Tubb4b | Hsp90b1 | Ccr2 | 1700020L24Rik |
| Ly6e | Rpl13a | Ubac1 | 1700020L24Rik | Prtn3 | Wfdc21 |
| Apoe | Rec114 | Col5a1 | Mpo | Mpo | Cebpe |
| Padi2 | Gata1 | Gfi1b | P4hb | Pld4 | Ltb4r1 |

| | | | | | |
|---|---|---|---|---|---|
| *Tbc1d4* | *Prss50* | *Tlk1* | *Pdia6* | *Gpr141* | *Trem3* |
| *Ier3* | *Rack1* | *Scrn3* | *Cebpe* | *F13a1* | *Lrg1* |
| *Stx3* | *Gclm* | *Slc43a1* | *Ctsg* | *Ly86* | *C3* |
| *Srgn* | *Slc25a21* | *Cd82* | *Rgcc* | *Tifab* | *B4galt6* |
| *Auts2* | *Slc45a3* | *Dut* | *Ly6c2* | *Itga1* | *Gda* |
| *Nfkbia* | *Rab4a* | *Acss1* | *Fkbp11* | *Emb* | *Hp* |
| *Itga2b* | *Rpl17* | *Fam210b* | *Sdf2l1* | *Ly6c2* | *Gfi1* |
| *Fut8* | *Rpl14* | *Zbtb46* | *Msrb1* | *Tspo* | *Prss57* |
| *Cpne2* | *Epdr1* | *Car1* | *Trem3* | *Mefv* | *Hdc* |
| *Mcpt8* | *Tnfaip2* | *Car2* | *Ap3s1* | *Trem3* | *Clec12a* |
| *Slc7a8* | *1700006J14 Rik* | *Larp1b* | *Ms4a3* | *Csf1r* | *Ms4a3* |
| *Ifitm1* | *Sdsl* | *Ppm1l* | *Pgam1* | *Unc93b1* | *Rgcc* |
| *Il1rl1* | *Il1rl1* | *Trim2* | *Ssr4* | *Ms4a4c* | *Mgst2* |
| *Runx1* | *Mrpl52* | *Hdgf* | *Atp8b4* | *Ms4a6c* | *Gstm1* |
| *Fyb* | *Prdx6* | *Lmna* | *Ssr2* | *Mpeg1* | *Pde4d* |
| *BC051537* | *Casp3* | *Dap3* | *Gsr* | *Pgam1* | *Gca* |
| *Csf2rb* | *Epor* | *Pklr* | *Tmed3* | *Cybb* | *Mrgpra2b* |
| *Ptprs* | *Clec4d* | *Selenbp1* | *Chst13* | *Cfp* | *Elane* |
| *Rab44* | *Mt1* | *Igsf3* | *Etfb* | *Gria3* | *Ctsg* |
| *H2-Oa* | *Aqp9* | *Gstm5* | *Lta4h* | *BC035044* | *Dstn* |
| *Casp3* | *Mns1* | *Gpsm2* | *Serpine2* | *Fcgr3* | *Nkg7* |
| *Galnt6* | *S100a10* | *Tmem56* | *Gpi1* | *Milr1* | *Prtn3* |
| *Tacstd2* | *Gnb4* | *Gclm* | *Selenom* | *Hexa* | *Ptgr1* |
| *Gng12* | *Plscr1* | *Fam241a* | *Ltb4r1* | *Mrpl33* | *Srgn* |

LT-HSC, long-term hematopoietic stem cell; ST-HSC, short-term hematopoietic stem cell; MPP, multipotent progenitor; MkP, megakaryocyte progenitor; BaP, basophil progenitor; CFU-E, colony forming unit - erythroid; GMP_mult, multipotent granulocyte macrophage progenitor; cMoP, common monocyte progenitor; GP, granulocyte progenitor.
